# Sfp1 integrates TORC1 and PKA activity towards yeast ribosome biogenesis

**DOI:** 10.1101/2022.12.02.518855

**Authors:** Luc-Alban Vuillemenot, Franz Y. Ho, Andreas Milias-Argeitis

## Abstract

Target of Rapamycin Complex 1 (TORC1) and Protein Kinase A (PKA) are two major regulators of cell growth in *Saccharomyces cerevisiae*, coupling nutrient availability with resource-intensive anabolic processes such as ribosome biogenesis. Even though TORC1 and PKA signaling converge on several shared targets, little is known on how these targets integrate signals from the two pathways. This is the case for Sfp1, a transcriptional activator of hundreds of ribosomal protein and ribosome biogenesis genes. To disentangle the roles of PKA and TORC1 in Sfp1 regulation, we constructed a large collection of Sfp1 (phospho)mutants and studied their intracellular localization and phosphorylation. Besides the already known TORC1 phosphorylation sites, we discovered that Sfp1 contains two PKA sites and a functional Nuclear Export Signal (NES) which is regulated by TORC1 and PKA. We further showed that TORC1 and PKA regulate Sfp1 phosphorylation independently of each other, with loss of activity of either pathway being sufficient to relocalize the protein from the nucleus to the cytoplasm, and the C-terminal zinc fingers being necessary for the responsiveness of Sfp1 to TORC1 and PKA inputs. This work contributes to our understanding of how cells regulate their growth by monitoring the outputs of multiple nutrient-sensing pathways.

## Introduction

Eukaryotic cells possess complex signaling networks which allow them to coordinate their growth with nutrient availability and the presence of stressors (Broach, 2012). In the model eukaryote *Saccharomyces cerevisiae* (budding yeast), the highly conserved Target of Rapamycin (TOR) and Protein Kinase A (PKA) kinases are two major regulators of cellular processes that govern many temporal and spatial aspects of cell growth (González and Hall, 2017; Wullschleger et al., 2006; Zaman et al., 2008; Zurita-Martinez and Cardenas, 2005). TOR functions within two distinct multiprotein complexes, TOR Complex 1 and 2 (TORC1 and TORC2), of which only TORC1 is rapamycin-sensitive (Loewith et al., 2002). TORC1 is known to promote resource-intensive anabolic processes such as ribosome biogenesis and protein translation while repressing catabolic processes such as autophagy, entry to G0 and the general stress response (Loewith and Hall, 2011; Wullschleger et al., 2006). Interestingly, PKA shares these functions with TORC1, as evidenced by the fact that TORC1 and PKA converge on a large set of common targets (Kunkel et al., 2019; Plank, 2022; Soulard et al., 2010), suggesting that cells integrate information from both pathways to make growth and developmental decisions. However, the two pathways have been traditionally studied separately from each other, and we still do not understand how joint TORC1 and PKA targets are regulated.

Ribosome biogenesis is a key cellular process that is jointly controlled by TORC1 and PKA in budding yeast, where the two pathways are known to regulate several key transcriptional activators (Sfp1, Ifh1) and repressors (Crf1, Tod6, Dot6, Stb3) (Huber et al., 2011; Lippman and Broach, 2009; Martin et al., 2004). Among the TORC1 and PKA targets in the ribosome biogenesis network, the *S*plit zinc-*F*inger *P*rotein 1 (Sfp1) stands out as a potent regulator of cell growth and cell size (Jorgensen et al., 2004, 2002). Sfp1 is a transcriptional activator implicated in the regulation of hundreds of growth-promoting genes, including most ribosomal biogenesis (Ribi), ribosomal protein (RP) and small nucleolar RNA (snoRNA) genes (Albert et al., 2019; Marion et al., 2004). *sfp1Δ* cells grow very slowly, they are the smallest of all single-deletion mutants with the same proliferation rate, and enter the cell cycle at a much smaller size compared to wild-type cells (Jorgensen et al., 2002). Under rich nutrient conditions Sfp1 localizes in the nucleus, but it rapidly relocalizes to the cytoplasm upon nutrient limitation or exposure to various stresses (Jorgensen et al., 2004; Lempiäinen et al., 2009; Singh and Tyers, 2009). Direct phosphorylation of Sfp1 by TORC1 at seven sites (Lempiäinen et al., 2009) (Fig. 1A) has been shown to regulate the subcellular localization of Sfp1, and TORC1 inhibition by rapamycin leads to a rapid exit of Sfp1 from the nucleus (Lempiäinen et al., 2009; Singh and Tyers, 2009). We have also recently shown that inhibition of PKA by the ATP analog 1-NM-PP1 in a background carrying analog-sensitive alleles of the PKA catalytic subunits, results in Sfp1 relocalization to the cytoplasm (Guerra et al., 2022), in agreement with earlier work suggesting that PKA regulates Sfp1 (Marion et al., 2004; Singh and Tyers, 2009). Still, it is unclear how this regulation is accomplished mechanistically. It has been shown that bovine PKA phosphorylates yeast Sfp1 *in vitro*, presumably through a conserved PKA consensus site (Budovskaya et al., 2005), but it remains unknown if budding yeast PKA phosphorylates Sfp1 *in vivo*. Moreover, joint regulation of Sfp1 activity by TORC1 and PKA raises the question of how TORC1- and PKA-dependent signals are integrated at the level of Sfp1.

**Figure 1:**
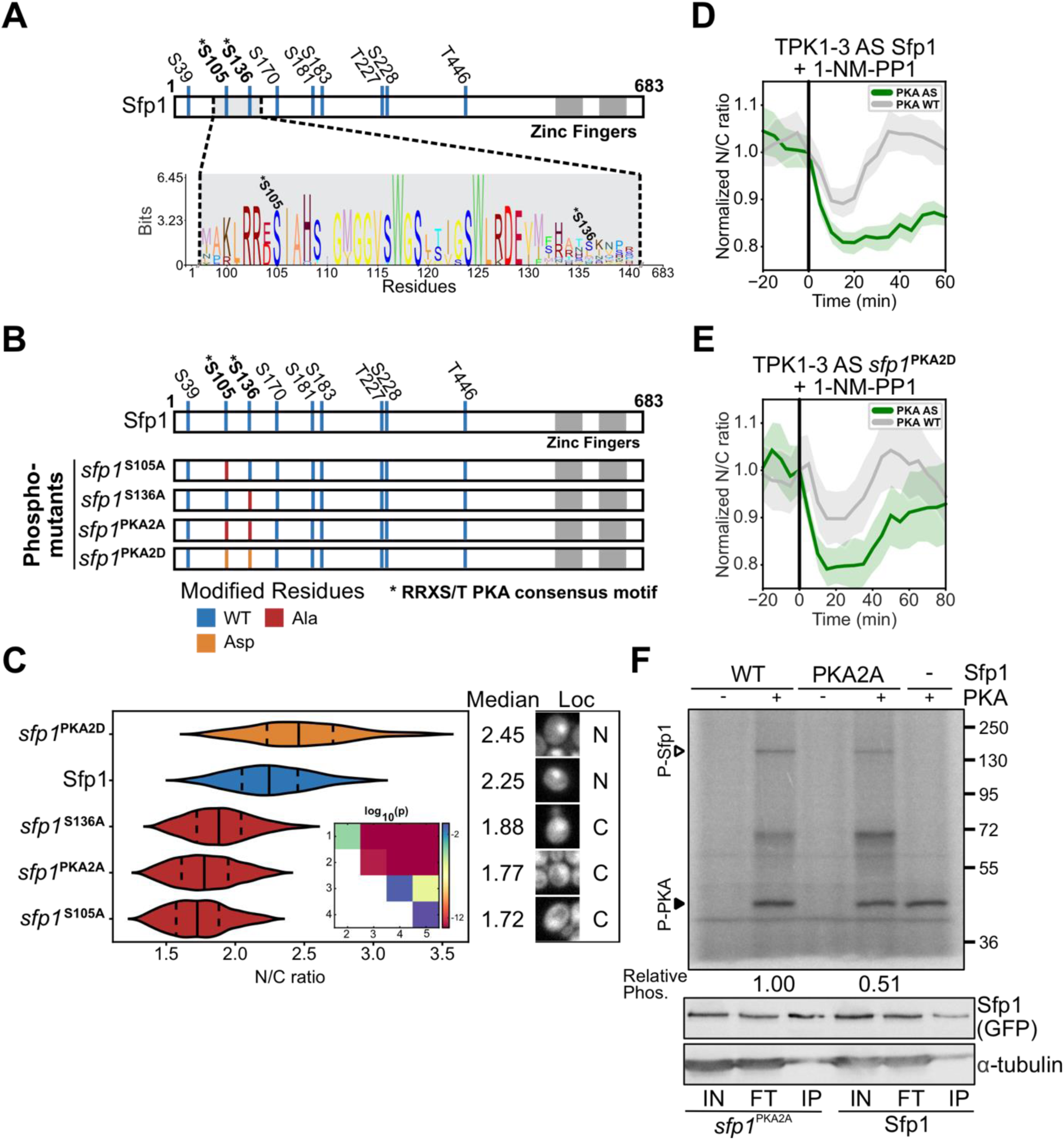
PKA directly regulates Sfp1 phosphorylation and localization. (**A**) Schematic representation of Sfp1. The TORC1 phosphorylation sites (blue marks, normal font), C-terminal Zn finger domains (dark grey segments) and the region containing the putative PKA sites (light grey segment, blue marks, bold font) are indicated. The sequence logos show the conservation of the highlighted fragment across different yeast genera (Fig.S1B), including the two putative PKA sites. (**B**) Scheme of wild-type Sfp1 and Sfp1 phospho-mutants carrying alanine (red marks) or aspartic acid (orange marks) substitutions at the PKA sites. (**C**) Sfp1 nuclear to cytosolic (N/C) ratio distributions in wild-type (n = 39) and cells carrying the *sfp1*^S105A^ (n = 50), *sfp1*^S136A^ (n = 41), *sfp1*^PKA2A^ (n = 46) and *sfp1*^PKA2D^ (n = 33) mutants. Violin plot colors follow the color code of panel B. Median (solid line) and 25th and 75th percentiles (dashed lines) are displayed for each distribution, and representative cell images are shown on the right. Statistical comparison of means was carried out with the Games-Howell test. P-values of all pairwise comparisons are graphically summarized in the inset matrix. Legend: 1 = *sfp1*^PKA2D^ 2 = Sfp1, 3 = *sfp1*^S136A^, 4 = *sfp1*^PKA2A^, 5 = *sfp1*^S105A^. Fig.S7A contains all p-values and 99% confidence intervals for all mean differences. (**D**) Sfp1 N/C ratio dynamics in response to 1-NM-PP1 addition in cells carrying the analog-sensitive PKA mutations (n = 50) compared to wild-type (n = 37). Off-target 1-NM-PP1 effects induce a transient drop in wild-type Sfp1 nuclear localization, whereas inhibition of PKAas leads to a sustained drop. In this and all following figures displaying N/C ratio dynamics, single-cell traces were normalized to the average N/C ratio at the moment of perturbation (t = 0, vertical line) in order to facilitate comparisons between mutants with different average N/C ratios. In all perturbation experiment figures, the average is displayed as a solid line and uncertainty bands represent the 95% confidence interval for the mean. (**E**) *sfp1*^PKA2D^ N/C ratio dynamics in response to 1-NM-PP1 addition in cells carrying the analog-sensitive PKA mutations (n = 27) compared to wild-type (n = 30). Inhibition of PKAas leads to a transient drop that is comparable to the 1-NM-PP1 off-target effect observed in wild-type *sfp1*^PKA2D^. The plot extends to 80 min post-treatment to demonstrate the convergence of the control and treated cells. Single-cell measurements at 65 min post-perturbation were discarded due to a transient shift in focus. Measurements at 60- and 70-min post-perturbation were linearly interpolated to generate the data point at 65 min. (**F**) Phosphorylation of Sfp1 by protein kinase A: following the treatment of PKAas cells with rapamycin and 1-NM-PP1 (to remove endogenous Sfp1 phosphorylation), Sfp1-pHtdGFP and *sfp1*^PKA2A^-pHtdGFP were immunoprecipitated by GFP-Trap (lower panel: Input (IN), Flow-through (FT), Immunoprecipitation (IP)), and were incubated with γ-^32^P-ATP, with or without bovine PKA for 30 min at 30°C (Upper panel). The reaction mixtures were separated by SDS-PAGE, the gel was exposed to storage phosphor screen, and autoradiography was performed by Cyclone plus (Perkin Elmer). The open triangle indicates the phosphorylated Sfp1 and the closed triangle indicates the (auto)phosphorylated PKA. Relative phosphorylation of Sfp1 in the two strains (Materials and Methods) is indicated below the autoradiograph. A replicate experiment is shown in Fig.S6.

To disentangle the roles of PKA and TORC1 in Sfp1 regulation, we generated a large collection of GFP-tagged Sfp1 (phospho)mutants, guided by a previous study of Sfp1 phosphorylation (Lempiäinen et al., 2009) and a sequence analysis of the protein which revealed the presence of two potential high-affinity PKA sites and a putative leucine-rich Nuclear Export Signal (NES). Through microscopic observation of the different Sfp1 mutants under steady-state growth and in response to dynamic perturbations of TORC1 and PKA activity, we demonstrated that Sfp1 contains at least two functional PKA phosphosites (verified by an *in vitro* kinase assay) and a functional NES whose activity is regulated by TORC1- and PKA-dependent phosphorylation. Using Phos-tag Western blotting of those mutants, we further showed that TORC1 and PKA regulate Sfp1 phosphorylation independently of each other, and that loss of phosphorylation from either pathway is sufficient to relocalize the protein from the nucleus to the cytoplasm. Therefore, our results suggest that high Sfp1 activity requires both TORC1 and PKA activity, and loss of input from either pathway is sufficient to downregulate Sfp1. Interestingly, although PKA seems to control a smaller number of phosphorylation sites compared to TORC1, it has an equally potent effect on Sfp1 localization. Finally, our results revealed that the C-terminal zinc fingers (Zn fingers) of Sfp1 are necessary for the dynamic response of its localization to perturbations of TORC1 and PKA activity, presumably via a binding partner.

Our work provides the first mechanistic demonstration of how a central regulator of ribosome biogenesis integrates inputs from the two major growth-regulatory pathways in budding yeast. We expect that similar mechanisms operate on other common TORC1 and PKA targets, for which Sfp1 can act as a prototype. Unraveling these mechanisms will be necessary for ultimately understanding how cells regulate their growth by monitoring and integrating the outputs of multiple nutrient-sensing pathways.

## Results

### PKA directly regulates Sfp1 phosphorylation and localization

Although it has long been known that Sfp1 localization depends on PKA activity levels (Marion et al., 2004; Singh and Tyers, 2009) and that bovine PKA phosphorylates Sfp1 *in vitro* (Budovskaya et al., 2005), it was unclear whether PKA phosphorylates Sfp1 *in vivo* and what is the effect of this phosphorylation. An analysis of the Sfp1 amino acid sequence revealed the presence of two high-affinity PKA RRxS consensus motifs (Plank et al., 2022), which we denoted by PKA1 (102-106, RRES*I) and PKA2 (133-137, RRNS*I). Alignment of Sfp1 homologues from different yeast genera with *Saccharomyces cerevisiae* Sfp1 (YLR403W) following the approach of (Pfanzagl et al., 2018) showed that PKA1 is highly conserved (Fig.1A). This observation prompted us to evaluate the functionality of PKA1 and PKA2 by constructing a set of phospho-mutants (Fig.1B). Our mutants contained alanine substitutions of PKA-targeted residues to prevent their phosphorylation, or mutations of these residues to aspartic acid to mimic phosphorylation. More specifically, we constructed *sfp1*^S105A^, *sfp1*^S136A^ and the double mutants *sfp1*^PKA2A^ (carrying both S105A and S136A) and *sfp1*^PKA2D^ (carrying S105D and S136D). These and all subsequently generated Sfp1 variants were C-terminally tagged with a pH-stable tandem GFP (Roberts et al., 2016) to enable their visualization with fluorescence microscopy. C-terminal tagging of Sfp1 has been used extensively in the past without adverse growth effects (Goranov et al., 2013; Guerra et al., 2022; Jorgensen et al., 2004; Lempiäinen et al., 2009; Singh and Tyers, 2009). However, due to the fact that some mutations could disrupt Sfp1 activity and disruption of Sfp1 leads to severe growth defects (Fingerman et al., 2003; Jorgensen et al., 2002), each Sfp1 mutant was genomically integrated in the HO locus (Voth et al., 2001) as an extra copy driven by the endogenous Sfp1 promoter, leaving the endogenous Sfp1 locus intact. Considering that this approach leads to an increase in overall Sfp1 levels in the cell and ectopic expression of Sfp1 reduces the nuclear localization of endogenous fluorescently-tagged Sfp1 (Singh and Tyers, 2009)(Fig.S1C), Sfp1-pHtdGFP was also integrated in the HO locus of the wild-type strain to provide the correct reference point for the nuclear localization of the Sfp1 mutants. To quantify the nuclear accumulation of Sfp1, all generated strains also carried a fluorescently tagged histone (Hta2-mRFP) which was used to locate the nucleus and quantify the nuclear-to-cytosolic (N/C) ratio of Sfp1 (Guerra et al., 2022). To facilitate data interpretation in the rest of the manuscript, it should be noted that, in our automated image analysis pipeline, an N/C ratio of around 1.5 corresponds to a largely cytosolic protein (cf. Materials and Methods).

Monitoring the single-cell N/C ratios of *sfp1*^S105A^ and *sfp1*^S136A^ under steady-state growth in glucose minimal medium (Methods), we observed that both mutations caused a massive relocalization of the protein to the cytoplasm compared to wild-type Sfp1 (Fig.1C), indicating that the PKA sites are functional under rich nutrient conditions, and are needed for keeping Sfp1 in the nucleus. Contrary to *sfp1*^PKA2A^ which was barely nuclear, the phosphomimetic *sfp1*^PKA2D^ was well-localized, and its N/C ratio was slightly higher than that of wild-type Sfp1. The fact that phosphomimetic mutations at the PKA-dependent sites promote Sfp1 localization while non-phosphorylatable residues delocalize the protein, suggested that the PKA sites of Sfp1 are functional.

To further support this hypothesis, we investigated the effect of acute PKA inhibition on the localization of wild-type Sfp1 and the PKA phosphomutants. To acutely inhibit PKA activity, we introduced the Sfp1 mutants in a PKAas strain background (Guerra et al., 2022), in which all three PKA catalytic subunits (Tpk1, Tpk2 and Tpk3) had been mutated to become sensitive to the ATP-competitive inhibitor 1-NM-PP1 (Zaman et al., 2009). Treatment of PKAas cells with 1-NM-PP1 led to a rapid and sustained drop in the Sfp1 N/C ratio (Fig.1D, Fig.S1D,E). On the other hand, the N/C ratio of *sfp1*^PKA2D^ in 1-NM-PP1-treated PKAas cells displayed only a transient drop that resembled the 1NM-PP1 control (Fig.1E, Fig.S1F,G). Moreover, no significant changes in localization were observed upon 1-NM-PP1 treatment of PKAas cells carrying *sfp1*^PKA2A^ (Fig.S1H,I), though this observation can also be attributed to the fact that this mutant was considerably cytosolic prior to 1-NM-PP1 addition. The divergent responses of wild-type Sfp1 and the PKA phosphomutants to 1-NM-PP1 further support the notion that PKA controls Sfp1 localization via the PKA1 and PKA2 sites.

To confirm that PKA is the kinase responsible for the phosphorylation of S105 and S136, we performed an *in vitro* kinase assay in which Sfp1-tdGFP and *sfp1*^PKA2A^-tdGFP purified from yeast were incubated with [γ-^32^P]ATP and the bovine PKA catalytic subunit (Materials and Methods). We observed that PKA can indeed phosphorylate Sfp1, and that PKA-dependent phosphorylation is reduced by 50% in *sfp1*^PKA2A^ (Fig.1F, replicate experiment Fig.S6). These results further support our notion that S105 and S136 are PKA-dependent phosphosites. They also suggest that Sfp1 likely contains additional PKA-dependent phosphosites which, as we will show below, are among the currently uncharacterized Sfp1 phosphosites.

Altogether, our observations under steady-state growth and upon PKA inhibition led us to conclude that PKA directly phosphorylates Sfp1 on the two identified consensus sites, and that PKA-dependent phosphorylation of those sites has a strong effect on Sfp1 localization.

### PKA and TORC1 regulate the nuclear export of Sfp1

Having established the presence and functionality of PKA sites on Sfp1, we next asked how PKA might regulate Sfp1 localization. A first hint was provided from the truncation of the N-terminal domain of Sfp1 (residues 1-202) which contains the two PKA and four TORC1 sites (Fig.2A). This truncation produced a mutant (Δ1-202, denoted by *sfp1*^ΔNter^) which was considerably more nuclear than wild-type Sfp1 (Fig.2C). Moreover, this mutant did not respond to rapamycin and methionine sulfoximine (MSX), a specific inhibitor of glutamine synthetase which perturbs TORC1 activity by depleting intracellular glutamine (Crespo et al., 2002; Guerra et al., 2022; Stracka et al., 2014) (Fig.S2A,B). These observations suggested that the N-terminal portion of Sfp1 contains a sequence that controls the nuclear exit of the protein.

**Figure 2:**
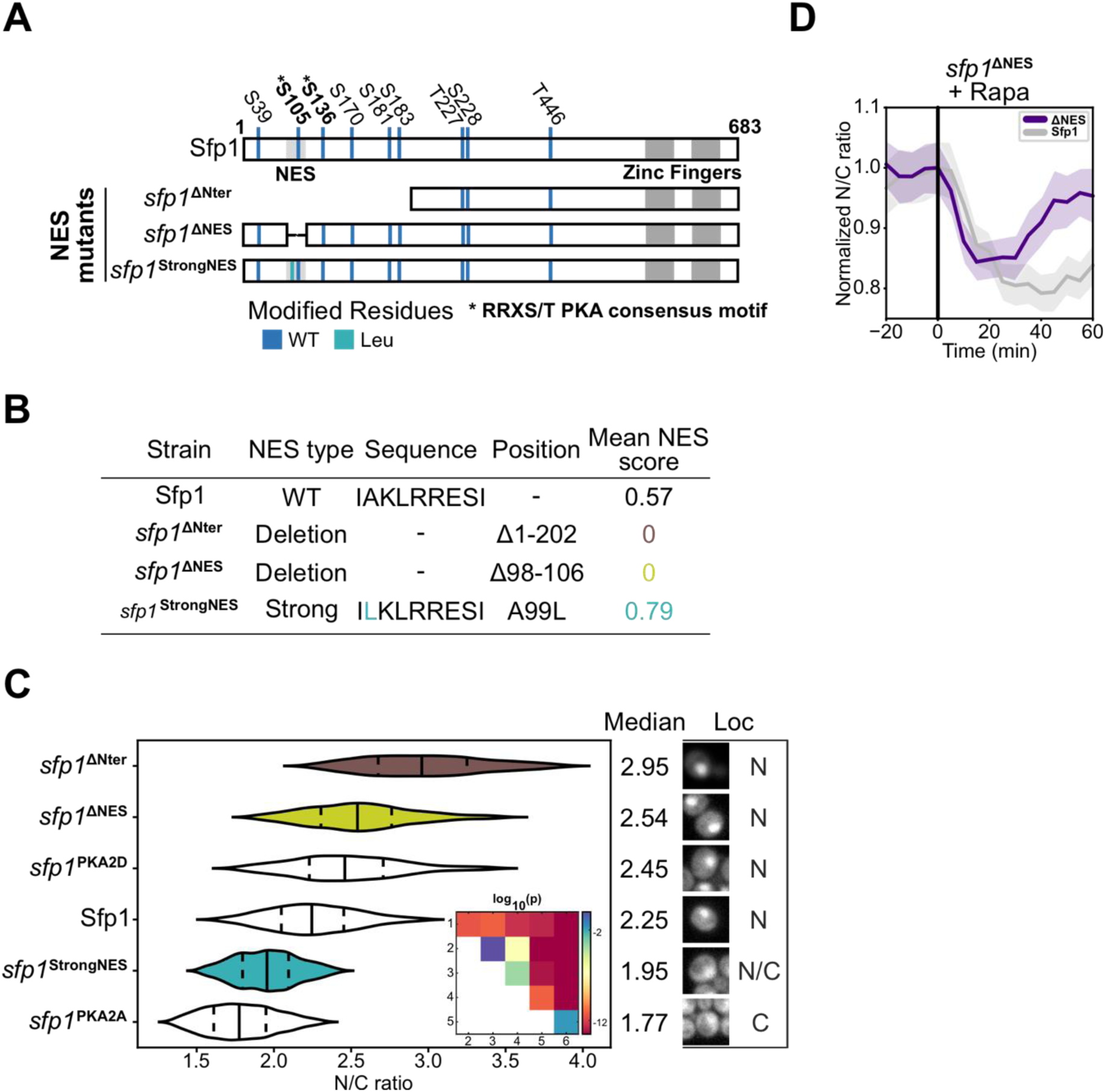
PKA and TORC1 regulate the nuclear export of Sfp1. (**A**) Schematic representation of mutants (deletion and point mutation) (**B**) Summary of NES scores for different Sfp1 mutants. NES scores were calculated with NetNES 1.1 (La Cour *et al*., 2004) (**C**) Sfp1 nuclear to cytosolic (N/C) ratio distributions in wild-type and cells carrying the *sfp1*^PKA2A^, *sfp1*^PKA2D^, *sfp1*^ΔNter^ (n = 49), *sfp1*^ΔNES^ (n = 30) and *sfp1*^StrongNES^ (n = 40) mutants. Data from wild-type, *sfp1*^PKA2A^ and *sfp1*^PKA2D^ same as in Fig. 1C, repeated here to facilitate comparisons. Median (solid line) and 25th and 75th percentiles (dashed lines) are displayed for each distribution, and representative cell images are shown on the right. Statistical comparison of means was carried out with the Games-Howell test. P-values of all pairwise comparisons are graphically summarized in the inset matrix. Legend: 1 = *sfp1*^ΔNter^ 2 = *sfp1*^ΔNES^, 3 = *sfp1*^PKA2D^, 4 = *sfp1*^StrongNES^, 5 = *sfp1*^PKA2A^. Fig.S7B contains all p-values and 99% confidence intervals for all mean differences. (**D**) Sfp1 N/C ratio dynamics in response to rapamycin in cells carrying wild-type Sfp1 and the *sfp1*^ΔNES^ mutant. Rapamycin leads to a sustained drop in wild-type Sfp1 nuclear localization (n = 47), whereas the *sfp1*^ΔNES^ mutant (n = 53) shows only a transient drop and recovers close to pre-treatment levels.

Closer examination of residues 1-202 with the NetNES prediction tool (La Cour et al., 2004) revealed the presence of a putative leucine-rich nuclear export signal (NES) at residues 98-106, overlapping with the PKA1 site and very close to PKA2 (Fig.2A). The location of the putative NES relative to the PKA sites led us to hypothesize that this NES is functional. To test this hypothesis, following the approach of (Ramakrishnan et al., 2019), we generated a potentially stronger NES by substituting an alanine with a leucine (A99L) in the identified sequence (Fig.2A). This change increased the leucine richness (and hydrophobicity) of the sequence, resulting in an increased *in silico* mean NES score (Fig.2B). In line with our expectations, the generated mutant (*sfp1*^StrongNES^) showed reduced nuclear localization compared to WT (Fig.2C), while still responding as expected to rapamycin and MSX treatments (Fig.S2C,D). In a second test, we deleted the putative NES sequence to generate *sfp1*^ΔNES^ (Δ98-106, Fig.2A,B). Similar to *sfp1*^ΔNter^, this mutant was more nuclear than the wild-type protein under steady-state conditions (Fig.2C). Overall, the *sfp1*^StrongNES^ and *sfp1*^ΔNter^ mutants produced phenotypes consistent with the presence of a functional NES.

The location of the NES relative to the PKA sites and our observations of *sfp1*^PKA2D^ and *sfp1*^PKA2A^ described above, strongly suggest that NES activity is inhibited by phosphorylation of the PKA1 and PKA2 sites: mimicking constitutive phosphorylation by PKA (*sfp1*^PKA2D^) results in a highly nuclear protein that responds to 1-NM-PP1 treatment only transiently (Fig.1E), whereas preventing the phosphorylation of PKA sites (*sfp1*^PKA2A^) causes the delocalization of the protein (active NES) from the nucleus. The proximity of several TORC1 sites to this NES (Fig.2A) further led us to hypothesize that NES activity is inhibited by phosphorylation of TORC1 sites in its vicinity. In support of this hypothesis, *sfp1*^ΔNES^ showed only a transient response to rapamycin, contrary to the sustained response of wt-Sfp1 (Fig.2D). Collectively, our observations suggest that 1) the identified putative NES is functional and 2) TORC1 and PKA phosphosites in the vicinity of this NES regulate its activity, and thus the exit rate of Sfp1 from the nucleus.

### PKA and TORC1 function independently on Sfp1

The evidence presented above suggests that PKA is a direct regulator of Sfp1. On the other hand, Sfp1 is a well-established TORC1 substrate (Lempiäinen et al., 2009). This fact raises the question of how Sfp1 integrates PKA and TORC1 inputs. To address the question, we generated a new set of phosphomutants (Fig.3A) and analyzed Sfp1 phosphorylation and localization using Phos-tag SDS-PAGE (O’Donoghue and Smolenski, 2022) and single-cell microscopy respectively. These analyses aimed to untangle the contributions of PKA and TORC1 on Sfp1 phosphorylation and understand how Sfp1 phosphorylation and localization are connected.

**Figure 3:**
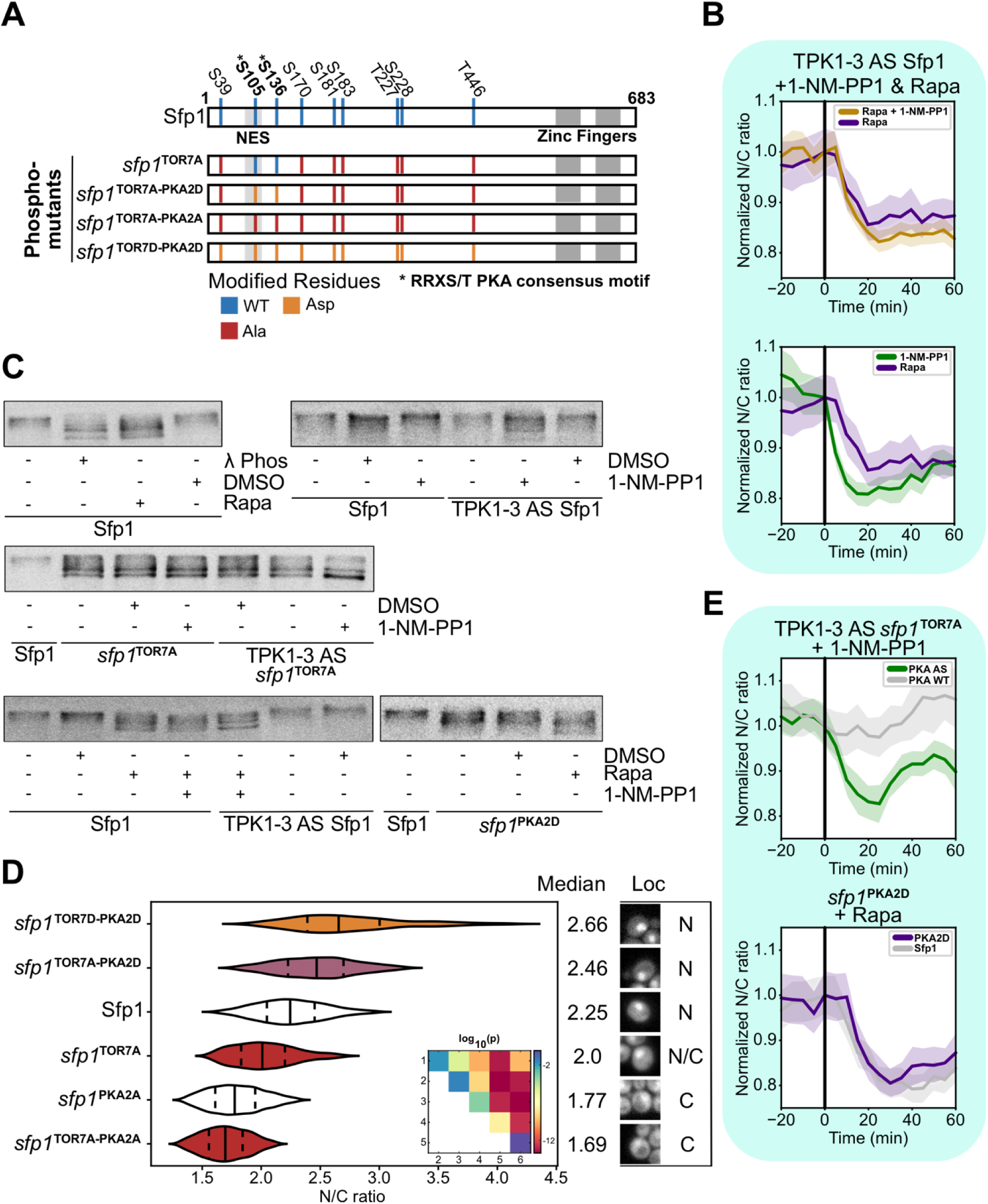
PKA and TORC1 function independently on Sfp1. (**A**) Schematic representation of Sfp1 phospho-mutants, following the color coding of Fig.1 (**B**) Response of Sfp1 to TORC1 inhibition by rapamycin (n = 40), PKA inhibition by 1-NM-PP1 (n = 50), and joint treatment with rapamycin and 1-NM-PP1 (n = 52), all in the PKAas background. (**C**) Phos-tag analyses of GFP-tagged Sfp1. λ phosphatase dephosphorylates serines, threonines and tyrosines. It was used to generate a (largely) dephosphorylated form of Sfp1 and facilitate visual comparisons with the phosphorylated forms. Vehicle (DMSO) and 1-NM-PP1 controls are also displayed to ensure that observed effects are due to inhibition of the respective kinases. (**D**) Sfp1 nuclear to cytosolic (N/C) ratio distributions in cells carrying the *sfp1*^TOR7D-PKA2D^ (n = 40), *sfp1*^TOR7A-PKA2D^ (n = 53), *sfp1*^TOR7A^ (n = 50) and *sfp1*^TOR7A-PKA2A^ (n = 33) mutants. Data from wild-type and *sfp1*^PKA2A^ same as in Fig. 1C, repeated here to facilitate comparisons. Median (solid line) and 25th and 75th percentiles (dashed lines) are displayed. Representative cells images are shown on the right of the violin plots. Statistical comparison of means was carried out with the Games-Howell test. p-values of all pairwise comparisons are graphically summarized in the inset matrix. Legend: 1 = *sfp1*^TOR7D-PKA2D^, 2 = *sfp1*^TOR7A-PKA2D^, 3 = Sfp1, 4 = *sfp1*^TOR7A^, 5 = *sfp1*^PKA2A^, 6 = *sfp1*^TOR7A-PKA2A^. Fig.S7C contains all p-values and 99% confidence intervals for all mean differences. (**E**) Sfp1 N/C ratio dynamics in cells carrying the *sfp1*^TOR7A^ (PKA AS n = 24, PKA WT n = 22) and *sfp1*^PKA2D^ (n = 33) mutants in response to 1-NM-PP1 and rapamycin respectively. *sfp1*^TOR7A^ still responds to 1-NM-PP1 inhibition of PKA, and *sfp1*^PKA2D^ still responds to rapamycin treatment.

Our first observation established that PKA and TORC1 have an equally strong impact on the localization of Sfp1, as the decrease in the Sfp1 N/C ratio was very similar between rapamycin and 1-NM-PP1 treatment of PKAas cells (Fig.3B). The fact that each pathway is capable of strongly de-localizing the protein is also reflected in the observation that simultaneous treatment of PKAas cells with rapamycin and 1-NM-PP1 does not further decrease the Sfp1 N/C ratio (Fig.3B, Fig.S3A).

Changes in Sfp1 localization follow changes in Sfp1 phosphorylation (Lempiäinen et al., 2009). However, it is not possible to infer the phosphorylation state of the protein based on its localization alone. For this reason, we employed a Phos-tag SDS-PAGE to assay Sfp1 phosphorylation under exponential growth in rich glucose medium and upon TORC1 and PKA inhibition (Methods). When cells were grown in YPD, Sfp1 produced a set of closely clustered bands in the Phos-tag gel (Fig.3C). Treatment with rapamycin resulted in a clear downward shift of those bands (Fig.3C), confirming that rapamycin leads to extensive dephosphorylation of the protein. Treatment of PKAas cells with 1-NM-PP1 produced only a subtle shift in the Sfp1 bands (Fig.3C), consistent with the fact that Sfp1 contains only a small number of PKA sites. Likely for the same reason (small number of PKA sites), double treatment with rapamycin and 1-NM-PP1 does not significantly decrease Sfp1 phosphorylation compared to rapamycin treatment alone (Fig.3C).

The fact that the overall phosphorylation of Sfp1 is only slightly reduced upon 1-NM-PP1 treatment suggests that the groups of PKA and TORC1-dependent phosphosites can be phosphorylated independently of each other. To further test this conjecture, we examined *sfp1*^TOR7A^, a phosphomutant which contains alanine substitutions on previously characterized TORC1 phosphosites (S39A, S170A, S181A, S183A, T227A, S228A & T446A, (Lempiäinen et al., 2009)) (Fig.3A). Due to the large loss of phosphorylation, this mutant produced bands that migrated further than wild-type Sfp1 (Fig.3C). However, 1-NM-PP1 treatment of PKAas cells carrying *sfp1*^TOR7A^ led to a further small reduction in phosphorylation (Fig.3C), suggesting that the PKA-dependent phosphosites are still phosphorylated in this mutant, despite the loss of seven TORC1 phosphosites. In a symmetrical fashion, rapamycin treatment of cells expressing *sfp1*^PKA2D^ led to a clear reduction in its phosphorylation (Fig.3C), and the same behavior was observed with *sfp1*^PKA2A^ (Fig.S3B). These observations suggest that the TORC1-dependent sites are functional independently of the status of PKA sites, and vice versa.

Our Phos-tag observations further suggested that the PKA and TORC1 phosphosites control Sfp1 localization independently of each other. To corroborate this hypothesis, we returned to microscopic observations of Sfp1 mutants (Fig.3D). *sfp1*^TOR7A^ appeared significantly (but not fully) delocalized, as had been observed before (Lempiäinen et al., 2009). However, 1-NM-PP1 treatment of PKAas cells carrying *sfp1*^TOR7A^ led to a further reduction in the N/C ratio of this mutant (Fig.3E, Fig.S3C), in line with the reduction in phosphorylation observed with the Phos-tag gel (Fig.3C,D). Testing the effect of rapamycin on *sfp1*^PKA2A^ was difficult since this mutant is almost fully delocalized, as discussed above. On the other hand, *sfp1*^PKA2D^ (which displayed good nuclear localization) still responded to rapamycin (Fig.3E), in agreement with the Phos-tag results which showed a large reduction in *sfp1*^PKA2D^ phosphorylation upon rapamycin treatment (Fig.3C).

To further test the independence of the TORC1 and PKA phosphosites, we generated phosphomutants that combined the alanine mutations at the TORC1 sites with alanine or aspartic acid mutations at the PKA sites (*sfp1*^TOR7A-PKA2A^ and *sfp1*^TOR7A-PKA2D^ respectively, Fig. 3A). *sfp1*^TOR7A-PKA2A^ (mimicking PKA inhibition in the *sfp1*^TOR7A^ background) was less localized compared to *sfp1*^TOR7A^ (Fig.3D), displaying one of the lowest N/C ratios observed in our experiments. On the other hand, the N/C ratio of *sfp1*^TOR7A-PKA2D^ (mimicking constitutive PKA phosphorylation) was greater than the N/C ratio of *sfp1*^TOR7A^ (Fig.3D). These results again highlight the fact the PKA phosphosites can alter protein localization independently of the TORC1 sites.

Taken together, our Phos-tag and microscopy data point to the fact that TORC1 and PKA control Sfp1 phosphorylation and localization independently of each other. Interestingly, although PKA controls a smaller fraction of the total Sfp1 phosphorylation compared to TORC1, it has an equally strong effect on Sfp1 localization.

### The C-terminal zinc fingers of Sfp1 are necessary for the dynamic response of Sfp1 localization

With PKA and TORC1 established as two major regulators of Sfp1 localization and phosphorylation, we reasoned that the localization of an Sfp1 mutant featuring phosphomimetic mutations on all known TORC1 and PKA residues (*sfp1*^TOR7D-PKA2D^) would display marginal or no sensitivity to chemical inhibition of TORC1 and PKA activity. To our surprise, and despite the fact that *sfp1*^TOR7D-PKA2D^ was highly nuclear as expected (Fig.3D), it was still clearly responsive to rapamycin, MSX and 1-NM-PP1 treatment (in PKAas cells) (Fig.4A, Fig.S4A,B), and so was *sfp1*^TOR7A-PKA2D^ (Fig.S4C-E). Previous work had suggested that there may be additional unmapped TORC1 sites on the protein (Lempiäinen et al., 2009), which we verified by observing that rapamycin treatment of *sfp1*^TOR7A^ led to a further decrease in its localization (Fig.S4F) and phosphorylation (Fig.S4G). We could also not exclude the presence of additional low-affinity PKA sites, or the possibility that TORC1 and/or PKA act upon Sfp1 both directly and via intermediate unidentified kinases.

**Figure 4:**
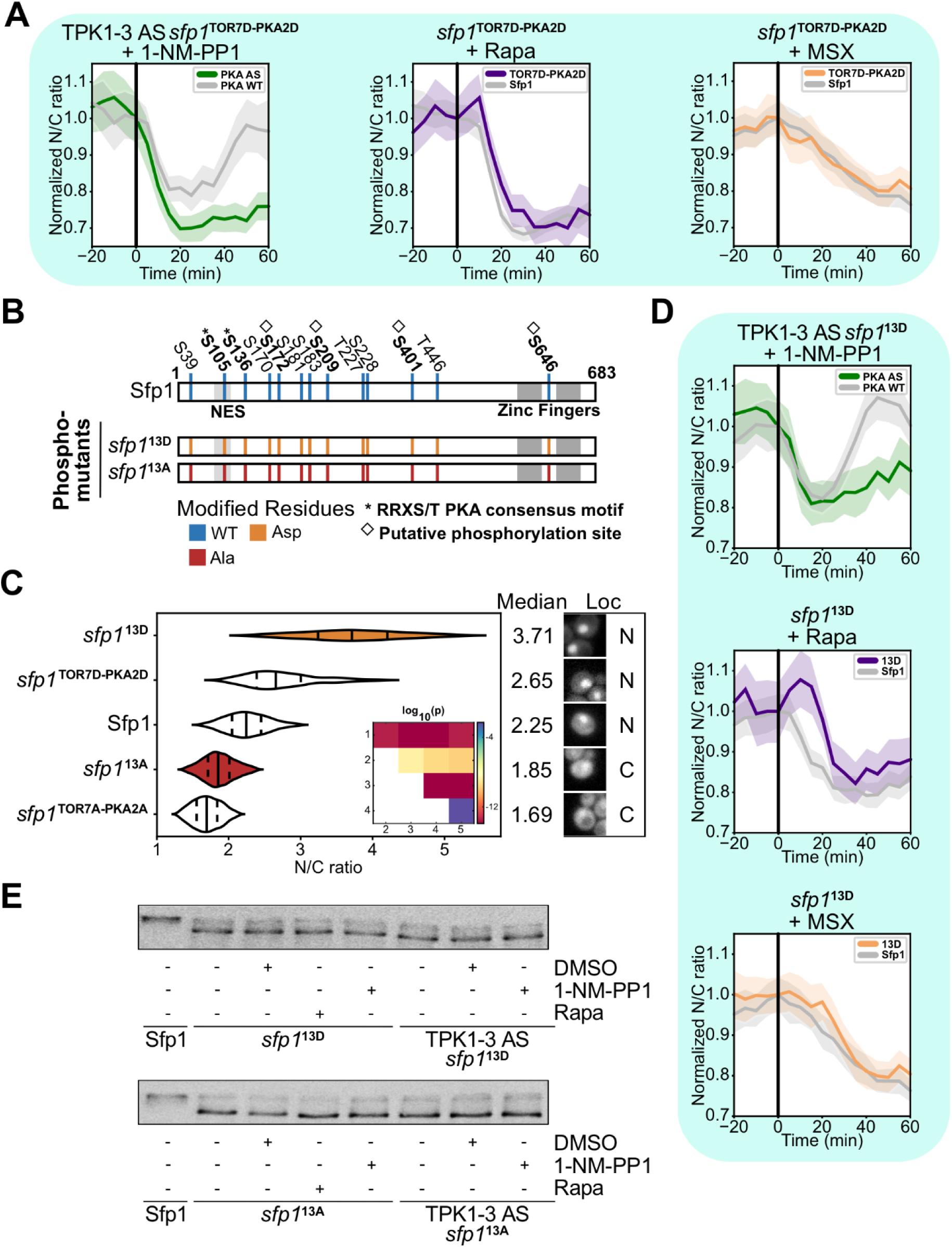
Another TORC1/PKA-dependent mechanism is involved in Sfp1 regulation. (**A**) Sfp1 N/C ratio dynamics in cells carrying the *sfp1*^TOR7D-PKA2D^ mutant in response to rapamycin (n = 27), methionine sulfoximine (n = 36) and 1-NM-PP1 (PKA AS n = 36, PKA WT n = 30). (**B**) Schematic representation of Sfp1 phospho-mutants, following the color coding of Fig.1. (**C**) Sfp1 nuclear to cytosolic (N/C) ratio distributions in cells carrying the wild-type, *sfp1*^13D^ (n = 33), *sfp1*^TOR7D-PKA2D^ (n = 40), *sfp1*^13A^ (n = 42) and *sfp1*^TOR7A-PKA2A^ mutants. Data from wild-type, *sfp1*^TOR7D-PKA2D^ and *sfp1*^TOR7A-PKA2A^ same as in Fig. 1C and Fig. 3D respectively, repeated here to facilitate comparisons. Median (solid line) and 25th and 75th percentiles (dashed lines) are displayed. Representative cells images are shown on the right of the violin plots. Statistical comparison of means was carried out with the Games-Howell test. P-values of all pairwise comparisons are graphically summarized in the inset matrix. Legend: 1 = *sfp1*^13D^, 2 = *sfp1*^TOR7D-PKA2D^, 3 = Sfp1, 4 = *sfp1*^13A^, 5 = *sfp1*^TOR7A-PKA2A^. Fig.S7D contains all p-values and 99% confidence intervals of all mean differences. (**D**) Sfp1 N/C ratio dynamics in cells carrying the *sfp1*^13D^ mutant in response to rapamycin (n = 35), MSX (n = 42) and 1-NM-PP1 (PKA AS n = 27, PKA WT n = 24). (**E**) Phos-tag analyses of GFP-tagged *sfp1*^13D^ and *sfp1*^13A^ in response to rapamycin and 1-NM-PP1 perturbations.

Recent work has uncovered several new phosphosites on Sfp1 (Lanz et al., 2021) which have not yet been linked to a known kinase. To test whether phosphomimetic mutations at those additional sites would result in a mutant that is completely insensitive to rapamycin and 1-NM-PP1, we generated *sfp1*^13D^, a mutant that carries four additional phosphomimetic substitutions (S172D, S209D, S401D, S646D) besides the ones of *sfp1*^TOR7D-PKA2D^ (Fig.4B). The large number of phosphomimetic mutations caused the N/C ratio of *sfp1*^13D^ to be the highest among all mutants that we generated (Fig.4C). Still, *sfp1*^13D^ responded to rapamycin, MSX and 1-NM-PP1 (in PKAas cells) (Fig.4D). This response suggested the presence of additional phosphosites on the protein. However, Phos-tag analyses of *sfp1*^13D^ (and *sfp1*^13A^) did not reveal visible changes in TORC1- or PKA-responsive phosphorylation on these mutants, which migrated as nearly single, unphosphorylated bands before and after treatment with rapamycin and 1-NM-PP1 (Fig.4E) and PKA-dependent phosphorylation of *sfp1*^13A^ was undetectable *in vitro* (Fig.S6). Therefore, if additional undiscovered TORC1-dependent phosphosites are present on the protein, they would have to be very few. An alternative (or additional) explanation for the response of the heavily modified *sfp1*^13D^ phosphomutant could be that an another TORC1- and/or PKA-sensitive mechanism is also implicated in the regulation of Sfp1 localization. Given that the C-terminal Zn fingers of Sfp1 are required for keeping the protein in the nucleus (Lempiäinen et al., 2009), we turned our attention to the C-terminus of Sfp1 for further clues.

Sfp1 is a zinc-finger protein, containing an N-terminal and two C-terminal Zn-finger domains of the C2H2 type, with the latter separated by an unusually long stretch of 40 amino acids. While the N-terminal Zn-finger does not seem critical for Sfp1 function (Fingerman et al., 2003), previous work had shown that disruption of the two C-terminal Zn-fingers greatly increases translation drug sensitivity of the resulting mutant and leads to loss of transcriptional activation of an Sfp1-responsive reporter (Fingerman et al., 2003). At the same time, disruption of these Zn-fingers leads to a reduction in the nuclear localization of the protein (Lempiäinen et al., 2009), suggesting that the Zn-fingers are crucial for proper Sfp1 localization and function.

To further explore the role of the C-terminal Zn-fingers in Sfp1 localization dynamics, we constructed *sfp1*^ΔCter^ (Fig.5A), a mutant in which the C-terminal part of Sfp1 encompassing the zinc finger domains was truncated (Δ490-683). Contrary to the N-terminally truncated *sfp1*^ΔNter^, (Fig.2C) GFP-tagged *sfp1*^ΔCter^ displayed a clear reduction in nuclear localization compared to wild-type Sfp1 (Fig.5B). Since the C-terminal truncation could affect localization by altering the structure of the protein, we also examined single and double Zn-finger mutants (Fig.5A), in which the conserved residues that typically chelate the zinc atom were converted to alanine (*sfp1*^Zn1^, *sfp1*^Zn2^ and *sfp1*^Zn3^; Zn1: C605A, H618A, H623A; Zn2: C661A, H664A, H677A, H680A; Zn3: combination of Zn1 and Zn2) (Fingerman et al., 2003; Lempiäinen et al., 2009). Consistently with previous work, the nuclear localization of these mutants was reduced compared to wild-type Sfp1 (Fig.5B). Moreover, *sfp1*^Zn3^ did not show any response to chemical perturbations that typically affect Sfp1 localization, such as rapamycin, MSX and 1-NM-PP1 treatment (Fig.5C, Fig.S5A,B), whereas mutants with a single Zn-finger disruption responded only very weakly (Fig.S5D,E). On the other hand, Phos-tag SDS-PAGE showed that the phosphorylation level of *sfp1*^Zn3^ was very close to wild-type, indicating that the large delocalization of the mutant was not a consequence of reduced phosphorylation. In line with this observation, the phosphorylation of *sfp1*^Zn3^ was still responsive to both rapamycin and 1-NM-PP1 (in PKAas cells) (Fig.5D), confirming that Zn-finger mutations do not prevent Sfp1 from being (de)phosphorylated, and suggesting that the Zn fingers are not essential for Sfp1 phosphorylation.

**Figure 5:**
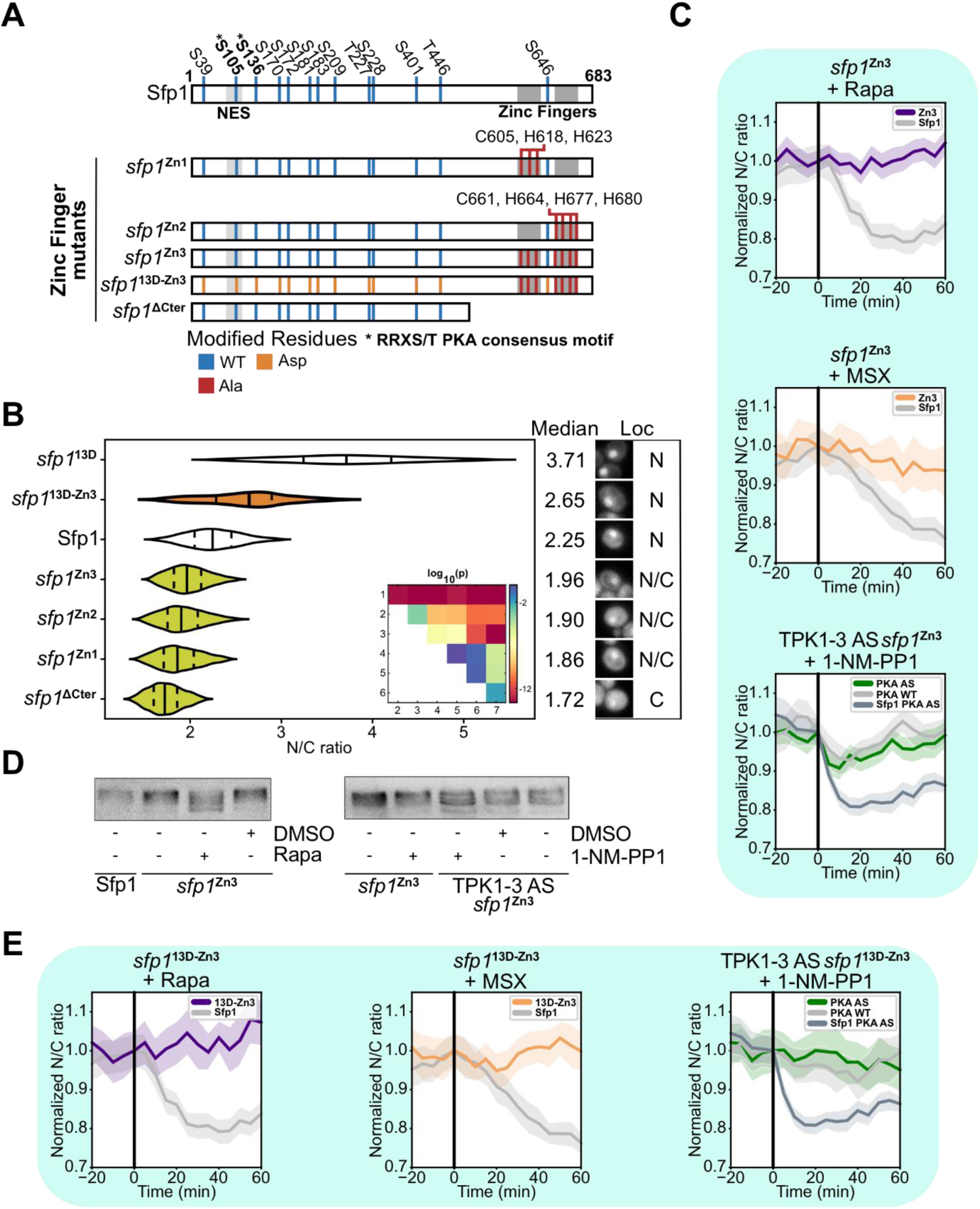
The C-terminal zinc fingers of Sfp1 are necessary for the dynamic response of Sfp1 localization. (**A**) Schematic representation of Sfp1 phospho- and Zn-finger mutants. (**B**) Sfp1 nuclear to cytosolic (N/C) ratio distributions in cells carrying the wild-type, *sfp1*^13D^, *sfp1*^13D-Zn3^ (n = 23), *sfp1*^Zn3^ (n = 45), *sfp1*^Zn2^ (n = 77), *sfp1*^Zn1^ (n = 91) and *sfp1*^ΔCter^ (n = 56) mutants. Data from wild-type and *sfp1*^13D^ same as in Fig. 1C and Fig. 4C respectively, repeated here to facilitate comparisons. Median (solid line) and 25th and 75th percentiles (dashed lines) are displayed. Representative cells images are shown on the right of the violin plots. Statistical comparison of means was carried out with the Games-Howell test. P-values of all pairwise comparisons are graphically summarized in the inset matrix. Legend: 1 = *sfp1*^13D^, 2 = *sfp1*^13D-Zn3^, 3 = Sfp1, 4 = *sfp1*^Zn3^, 5 = *sfp1*^Zn2^, 6 = *sfp1*^Zn1^, 7 = *sfp1*^ΔCter^. Fig.S7E contains all p-values and 99% confidence intervals of all mean differences. (**C**) Sfp1 N/C ratio dynamics in cells carrying the *sfp1*^Zn3^ mutant in response to rapamycin (n = 56), MSX (n = 26) and 1-NM-PP1 (PKA AS n = 46, PKA WT n = 37). (**D**) Phos-tag analyses of GFP-tagged *sfp1*^Zn3^ in response to rapamycin and 1-NM-PP1 perturbations. (**E**) Sfp1 N/C ratio dynamics in cells carrying the *sfp1*^13D-Zn3^ mutant in response to rapamycin (n = 32), MSX (n = 35) and 1-NM-PP1 (PKA AS n = 37, PKA WT n = 35).

Given that *sfp1*^Zn3^ displays very low nuclear localization, the lack of response to chemical inhibitors of TORC1 and PKA (Fig.5C) could be due to the fact that further decreases in the N/C ratio would be hard to quantify. We therefore inserted the Zn-finger mutations to our phosphomimetic mutants, since we had observed that phosphomimetic mutations result in increased Sfp1 localization (Fig.3D, Fig. 4C). In line with our expectations, all “hybrid” mutants combining phosphomimetic substitutions with Zn-finger mutations (*sfp1*^TOR7D-PKA2D-Zn1^, *sfp1*^TOR7D-PKA2D-Zn2^, *sfp1*^TOR7D-PKA2D-Zn3^, *sfp1*^13D-Zn3^) were clearly nuclear (Fig.5B, Fig.S5C). Surprisingly, however, mutation of the Zn-fingers in the phosphomimetic *sfp1*^13D^ protein had an unexpected effect: in contrast to *sfp1*^13D^ which responded to rapamycin and MSX treatment (Fig.4D), the corresponding Zn-finger mutant (*sfp1*^13D-Zn3^, Fig.5A) did not (Fig.5E, Fig.S5H,I). Moreover, the same behavior was observed with *sfp1*^TOR7D-PKA2D-Zn1^, *sfp1*^TOR7D-PKA2D-Zn2^ and *sfp1*^TOR7D-PKA2D-Zn3^ (Fig.S5F,G). In brief, introduction of Zn-finger mutations to Sfp1 phosphomutants made the resulting hybrid mutants unresponsive to TORC1 and PKA perturbations.

Taken together, our observations of the Zn-finger mutants suggest that, besides the NES at the N-terminus of the protein, the C-terminal Zn fingers of Sfp1 are essential for the response of Sfp1 to changes in TORC1 and PKA activity. This C-terminal mechanism does not seem to interfere with the phosphorylation of TORC1- and PKA-dependent sites, and its precise nature remains elusive (some possible alternatives are provided in the Discussion).

## Discussion

Through microscopic observations and phosphorylation analyses of a large collection of Sfp1 mutants, we found that Sfp1 integrates signals from both TORC1 and PKA pathways. Specifically, Sfp1 contains at least two functional PKA phosphorylation sites which, together with TORC1 phosphosites, regulate the activity of an N-terminal nuclear export sequence. Our *in vitro* kinase activity assays with PKA suggest that Sfp1 contains additional phosphosites besides the ones determined here. Given the fact that *sfp1*^13A^ does not show any PKA-dependent phosphorylation (Fig.S6), the additional PKA-dependent sites are very likely among the sites that are mutated in *sfp1*^13A^.

Loss of activity from either pathway (TORC1 or PKA) is sufficient to delocalize Sfp1 via loss of phosphorylation at the corresponding sites, suggesting that nuclear accumulation of Sfp1 requires both TORC1 and PKA activity. Interestingly, translocation of Sfp1 into the nucleus appears to be regulated by an additional mechanism embedded in the C-terminal domain of the protein, and intact Zn fingers are a prerequisite for the response of Sfp1 to the loss of TORC1 and PKA activity. Given that phosphomimetic mutations can restore the localization of Zn finger mutants and that Zn finger mutants are highly phosphorylated, the N-terminal (phosphorylation-based) and C-terminal (Zn finger-based) mechanisms seem to operate independently of each other.

The implication of the C-terminal Zn fingers in the translocation of Sfp1 is surprising, given that Zn fingers are typically considered to be DNA-binding domains. However, C2H2 Zn fingers such as those of Sfp1 can also mediate protein-protein interactions (Brayer and Segal, 2008), and prior work has provided evidence that the Rab escort protein Mrs6 is implicated in the regulation of the Sfp1 localization (Lempiäinen et al., 2009; Singh and Tyers, 2009). Intriguingly, the Zn1 and Zn2 mutants show a severe interaction defect with Mrs6 (Lempiäinen et al., 2009), which suggests that the C-terminal Zn fingers mediate the interaction of the two proteins. However, experimental evidence on the role of Mrs6 is contradictory: it has been suggested that Mrs6 acts as a nutrient-sensitive cytoplasmic anchor that only interacts with Sfp1 under poor nutrient conditions (Singh and Tyers, 2009), or that it promotes Sfp1 nuclear localization and phosphorylation by interacting with Sfp1 in rich nutrients (Lempiäinen et al., 2009). One cannot exclude the possibility that Mrs6 itself receives signals from TORC1 and/or PKA, which in turn cause changes in the Sfp1-Mrs6 interaction and provide a second translocation mechanism besides the N-terminal mechanism described here. Yet, no TORC1- or PKA-dependent input signals for Mrs6 are known to date. Studies of the Mrs6-Sfp1 interaction are complicated by the fact that Mrs6 is an essential Rab escort protein involved in intracellular trafficking (Fujimura et al., 1994; Singh and Tyers, 2009), whose mutation or overexpression can readily alter cell physiology. Although Mrs6 is a promising candidate for mediating changes in Sfp1 localization, it should be noted that the C-terminal translocation mechanism of Sfp1could also operate without Mrs6. For example, a Nuclear Localization Signal (NLS) could be embedded in the Zn finger region of Sfp1, since DNA-binding regions and NLSs frequently overlap (Cokol et al., 2000). The activity of such an NLS could be further modulated by phosphorylation of a few still-uncharacterized TORC1- and PKA-dependent phosphosites closer to the C-terminus. Structural analysis of Sfp1 could also determine if the C- and N-termini of the protein interact, connecting changes in N-terminal phosphorylation into changes in DNA binding and localization. Overall, the C-terminal regulation of Sfp1 localization warrants further study.

In summary, our work demonstrates that Sfp1 integrates inputs from TORC1 and PKA, the two major growth-regulatory pathways of budding yeast. While the precise relation of the TORC1 and PKA pathways and their crosstalk are still being debated (Plank, 2022), our results show that TORC1 and PKA function independently of each other towards Sfp1, and loss of input from either pathway results in the relocalization of Sfp1 to the cytoplasm and a subsequent decrease in Sfp1 activity. By using two independent inputs, cells are able to adjust Sfp1 activity in response to different external and internal metabolites sensed by TORC1 and PKA, and ensure that Sfp1 is maximally active only when both signaling pathways are active. Future studies of other major TORC1 and PKA targets will be necessary for understanding how cells make decisions by integrating information from these central growth regulators.

## Materials and Methods

### Plasmid construction

A pH-resistant tandem GFP (pHtdGFP, (Roberts et al., 2016)) was used to C-terminally tag all Sfp1 mutants generated in this work, with a short linker (GDGAGLIN) inserted between GFP and each Sfp1 mutant. Sfp1 mutations were carried out by site-directed mutagenesis with PCR amplification using Q5 & Phusion polymerases (New England Biolabs). For plasmids pLV30 (*sfp1*^Zn1^), pLV31 (*sfp1*^Zn2^) & pLV58 (*sfp1*^TOR7A^), Sfp1 mutants were linearized from the yeast strains HL215, HL350 & HL350 (Lempiäinen et al., 2009). DNA synthesis (Twist Bioscience) was used to generate the *sfp1*^13D^ and *sfp1*^13A^ mutants. Plasmids carrying the Sfp1-pHtdGFP mutants were built via Gibson Assembly and propagated in *E. coli* DH5α cells. All plasmids were verified by Sanger sequencing (Eurofins genomics). Further details about the plasmids used in this study can be found in Table S1.

### Yeast strains

The *Saccharomyces cerevisiae* S288C-derived prototrophic YSBN6 background strain (Canelas et al., 2010) was used to construct all the yeast strains presented in this study. Plasmids were linearized by PCR for stable genomic integration in yeast using the classical Lithium Acetate transformation (Gietz and Schiestl, 2007). Sfp1 mutants driven by the Sfp1 promoter were integrated by homologous recombination in the HO locus of YSBN6 cells, leaving the endogenous Sfp1 locus intact. Additionally, all strains expressed an mRFP-tagged histone (Hta2) as a nuclear marker. Each genomic construct was verified by PCR and Sanger sequencing (Eurofins genomics, The Netherlands). A list of all the yeast strains generated in this study can be found in Table S2.

### Growth conditions

For all microscopy experiments, cells were grown in minimal medium (Verduyn et al., 1992) supplemented with 2% glucose (Sigma-Aldrich). Prior to microscopy experiments, cells were grown overnight in batch culture at 30°C with shaking at 300 rpm, diluted the next morning and grown until exponential phase. For all Western blot experiments, cells were grown in YPD (Glucose 2%) under the same batch culture conditions (30°C with shaking at 300 rpm, grown overnight and diluted in the next morning).

### Drug preparations

To avoid a strong effect of Dimethyl sulfoxide (DMSO) on the localization of Sfp1-pHtdGFP (and its variants), Rapamycin (Sigma-Aldrich) was diluted in minimal medium for microscopy experiments. Rapamycin powder was dissolved in pure methanol to prepare a concentrated stock at 30 µg/mL which was sterilely aliquoted in Eppendorf tubes to let methanol evaporate prior to storage at -80°C. For each chemical perturbation experiment, a dried rapamycin aliquot was taken out of the freezer to be resuspended in pre-warmed minimal medium (supplemented with 2% glucose). The acute inhibition of TORC1 was achieved by addition of 2 µL of the medium-resuspended rapamycin solution (200 ng/mL final concentration) directly to the cells in the plastic chambered coverslip. For the western blot experiments, rapamycin was diluted in DMSO and delivered to the batch cultures at a final concentration of 200 ng/mL. Methionine Sulfoximine (MSX, Sigma-Aldrich) was diluted in H_2_O for microscopy experiments and used at final concentration of 0.36 mg/mL. 1-NM-PP1 (Sigma-Aldrich) was a diluted in DMSO at a final concentration of 0.5µM for microscopy and Western blot experiments.

### Microscopy

Inverted widefield fluorescence microscopes (Eclipse Ti-E, Nikon instruments), fitted with microscope incubators (Life Imaging Services), were used to carry out all microscopy experiments. A constant temperature of 30°C was maintained over the span of each experiment. 100x Nikon S Fluor Iris (NA=1.30), 100x Nikon Apo λ Oil (NA=1.45) or 100x Nikon Apo VC Oil DIC N2 (NA=1.40) objectives were used for the experiments. To record the microscopy images, iXon Ultra 897 DU-897-U-CD0-#EX cameras (Andor Technology) were used. The Perfect Focus System (PFS) was used to prevent loss of focus during the experiments. Two different LED-based excitation systems were used to carry out fluorescence measurements: pE2 (CoolLED Limited) and Lumencor (AURA). For GFP imaging, cells were excited at 470 nm (excitation filter, 450-490 nm; dichroic, 495 nm; emission filter, 500-550 nm) while RFP imaging was carried out by exciting cells at 565 nm (excitation filter, 540-580 nm; dichroic, 590 nm; emission filter, 600-650nm). During brightfield imaging, a long-pass filter (600 nm) was used.

To determine the nuclear-to-cytosolic ratio of the different Sfp1 mutants during steady-state growth, exponentially growing cells, cultivated at 30°C in minimal medium supplemented with 2% glucose, were inoculated under prewarmed minimal medium agar pads (2% glucose, 1% agarose), subsequently incubated at 30°C for half an hour under the microscope, and then imaged for at least 12.5 hours. For these experiments, several non-overlapping xy positions were recorded, with brightfield, GFP and RFP images acquired at each position every 5 min for the whole duration of the experiment.

To follow changes in the nuclear-to-cytosolic ratio of Sfp1 in response to chemical inhibition of TORC1 and PKA, plastic chambered coverslips (µ-Slide 8 Well, Ibidi) were coated with Concanavalin A (Sigma-Aldrich) at a final concentration of 1 mg/mL. Specifically, each well was loaded with 250 µL of Concanavalin incubated for 10 min in a sterile environment, washed with 250µL sterile ddH_2_O and dried for 20 min. 300µL of exponentially growing cells, cultivated in minimal medium (supplemented with 2% glucose) at 30°C, were loaded at an Optical Density (OD) at 600 nm of 0.1 and incubated for 30 min at 30°C. To remove unattached cells, the wells were then washed twice with pre-warmed minimal medium (supplemented with 2% glucose). The attached cells were subsequently imaged under the microscope in the brightfield, GFP and RFP channels with the imaging settings described above. Images were recorded every 5 min at multiple non-overlapping xy positions for each well. Cells were treated with the following chemicals: the drug vehicle DMSO (0.16 % final concentration, Sigma-Aldrich), 1-NM-PP1 (final concentration 0.5 µM, diluted in DMSO), Rapamycin (final concentration 200 ng/mL, diluted in minimal medium) and Methionine Sulfoximine (MSX; final concentration 0.36 mg/mL, diluted in H_2_O). The chemicals were pipetted into the wells between two recordings, approximately after 65 min after the start of the experiment.

### Image analysis and calculation of Sfp1-pHtdGFP single cell Nuclear-to-Cytosolic ratios

For steady-state and chemical perturbation experiments, the rolling ball background subtraction plugin of ImageJ (v.1.52n, Java 1.8.0_202) was used for background correction of fluorescence channel images (GFP & RFP). Mother cells were segmented and tracked in brightfield using the semi-automatic ImageJ plugin BudJ (Ferrezuelo et al., 2012). All strains expressing Sfp1-pHtdGFP mutants also expressed an RFP-tagged histone (Hta2-mRFP) as a nuclear marker. For each segmented cell, a smaller mask corresponding to the nucleus was determined based on Hta2-mRFP (for further details, see (Guerra et al., 2022)). The cell and nuclear masks were used to obtain the average GFP pixel intensity in the nucleus and cytoplasm in order calculate the GFP nuclear-to-cytosolic (N/C) ratios for individual cells using the custom-made Python script that was presented in (Guerra et al., 2022) and is available online (https://github.com/amiliasargeitis/microscopy_scripts/). For each time-lapse microscopy experiment involving chemical inhibition of TORC1 and/or PKA, single-cell time traces of (mutant) Sfp1 N/C ratios were divided by the corresponding population average of the N/C ratio at the moment of treatment. In this way, all single-cell time series were normalized to a mean of 1 at the moment of treatment, to facilitate comparisons across mutants with different steady-state N/C ratios. Our plots report the averages of single-cell N/C ratio traces (solid lines), along with the 95% confidence intervals of the mean (shaded bands). Microscopic quantification of (mutant) Sfp1 localization during steady-state growth followed the same analysis pipeline with the chemical perturbation experiments with the exception of normalization. Before constructing the violin plots, the upper and lower 2.5% of the single-cell N/C ratios (typically corresponding to outliers) were removed from the dataset of each strain.

### SDS-PAGE Phos-tag and western Blotting

To monitor changes in the phosphorylation of our GFP-tagged Sfp1 mutants, exponentially growing YPD cultures (OD_600nm_ = 0.8) were treated with rapamycin (final concentration: 200 ng/mL, diluted in DMSO), 1-NM-PP1 (final concentration: 0.5 µM, diluted in DMSO) or DMSO (Drug vehicle controls were carried out with the same DMSO volume used for the drug treatments), and sampled 30 min after treatment. Following the addition of inhibitors (for details, see Drug Preparations), 10 mL of exponentially growing cells were sampled. The samples were fixed with cold Trichloroacetic Acid (final concentration 6%), kept in ice for 10 min, pelleted, washed twice with cold acetone and air-dried. The pellets were resuspended in urea buffer (50 mM Tris HCl pH 7.5, 6 M Urea and 1% Sodium Dodecyl Sulphate supplemented with 100mM phenylmethylsulfonyl fluoride and 1% phosphatase inhibitor cocktail 3 (Sigma-Aldrich)), and cell lysis was carried out by 10 min-long bead beating. To precipitate insoluble proteins, the samples were incubated for 10 min at 65°C (shaking at 800 rpm) and centrifuged (15000 rpm) at 4°C for 5 min. The supernatant was transferred in new Eppendorf tubes, protein concentration was measured with a Nanodrop spectrophotometer (Nanodrop 2000, Thermo Fisher) and the samples were then diluted. To determine the dephosphorylated state of Sfp1, the wild-type protein extracts were incubated with 40 units of Lambda Protein Phosphatase (New England Biolabs, P0753S) at 30°C for 30min. For each Phos-tag Western blot experiment, proteins denatured in SDS sample buffer [50mM Tris HCl pH 6.8, 2% (w/v) SDS, 1.075 M Glycerol, 1.5mM Bromophenol blue, 1% (w/v) 2-Mercaptoethanol], were resolved on 7.5% polyacrylamide gels [0.375 M Tris, 50 µM Phos-tag (Fujifilm Wako Chemicals), 0.1 mM MnCl_2_], subsequently transferred to polyvinylidene fluoride membranes and probed with the anti-GFP primary antibody (Roche, Mouse monoclonal, #11814460001) and goat AffiniPure anti-mouse IgG (H+L) HRP conjugate secondary antibody (Jackson ImmunoResearch, Goat polyclonal, #115-035-003).

### *In vitro* kinase assay

Exponentially growing cells expressing wild type Sfp1, *sfp1*^PK2A^ and *sfp1*^13A^ (in PKAas background) were cultured in rich glucose medium (YPD; 50 mL) and, prior to cell lysis, were subjected to a 30-min treatment with rapamycin (final concentration: 200 ng/mL, diluted in DMSO) and 1-NM-PP1 (final concentration: 0.5 µM, diluted in DMSO) to ensure dephosphorylation of the protein. Cells were collected at OD_600_ = 1, washed once at 4°C in cold ddH2O, resuspended in lysis buffer (10 mM Tris/Cl pH 7.5, 150 mM NaCl, 0.5 mM EDTA, 0.5 % Nonidet^TM^ P40 Substitute 100mM phenylmethylsulfonyl fluoride and 1% protease inhibitor cocktail (Sigma-Aldrich)), and cell lysis was carried out by 10 min-long bead beating. To precipitate insoluble proteins the samples were incubated centrifuged (15000 rpm) at 4°C for 5 min and the supernatant was transferred in new Eppendorf tubes. The proteins were immunoprecipitated using GFP-Trap (Chromotek) according to manufacturer’s protocol (with slight modifications). To remove any remaining phosphorylation the GFP-Trap bound proteins were treated with 40 units of Lambda Protein Phosphatase (New England Biolabs, P0753S) at 30°C for 30min. The GFP-Trap bound proteins were then washed once with kinase buffer (25 mM Tris (pH 7.5), 130 mM NaCl, 15 mM MgCl_2_, 20 mM β-Glycerol phosphate, 1 mM NaF, 1 mM Na_3_VO_4_ and 2 mM DTT) and resuspended in 40 µl kinase buffer containing 300 µM ATP with 6 µCi γ-^32^P-ATP (Hartmann analytics) and 10 U bovine PKA (Sigma, 539576). For no PKA control, kinase buffer was added instead. The reaction mixtures were incubated in a 30°C heat block for 30 min with manual occasional mixing, and then quenched by adding 5X Laemmli buffer (Laemmli UK,1970) to the final concentration of 1X. The reaction mixtures were heated at 95 °C for 10 min and 25 µl of each samples were separated by 9% SDS-PAGE. After electrophoresis, the gel was fixed in 40% ethanol, 10% acetic acid for 15 min, rinsed with water and stained by InstantBlue Coomassie Protein Stain (Abcam). The stained gel was then rinsed again with water and dried by a Gel Dryer (Bio-Rad). The dried gel was then exposed to a Storage Phosphor Screen (Packard) and the screen was scanned by Cyclone Plus (Perkin Elmer).

### Statistical comparisons

All statistical comparisons of average N/C ratios were carried out with the R implementation of the Games-Howell non-parametric multiple comparisons test (Zar, 1999) from the `rstatix` package (https://rpkgs.datanovia.com/rstatix/reference/games_howell_test.html). A multiple comparisons test is necessary to avoid inflating the probability of a Type I error that would otherwise result from multiple pairwise comparisons using a standard t-test. The Games-Howell test in particular is applicable when the variances of the compared populations are unequal, as is the case with our Sfp1 N/C ratio data (Zar, 1999).

### Sfp1 sequence analysis

Alignment of the homologous sequences to *S. cerevisiae* Sfp1 was carried out similarly to (Pfanzagl et al., 2018). Specifically, 22 Sfp1 and Sfp1-like proteins across several yeast genera were aligned with the MUSCLE (Multiple Sequence Comparison by Log-Expectation) algorithm (cluster method: neighbour joining) and visualized using Jalview (Fig. S1). The multiple sequence alignment was then used to generate the Hidden Markov Model (HMM) profile by using Skylign (http://skylign.org, (Wheeler et al., 2014)).

## Acknowledgements

We would like to thank David Shore (University of Geneva, Switzerland) for providing strains from (Lempiäinen et al., 2009).

## Author Contributions

Conceptualization: L-A.V., A.M.-A.; Methodology: L-A.V., A.M-A.; Software: L-A.V.; Formal analysis: L-A.V.; Investigation: L-A.V., F.H.Y.; Resources: L-A.V., F.Y.H.; Data curation: L-A.V.; Writing -original draft: L-A.V., A.M.-A.; Visualization: L-A.V.; Supervision: A.M.-A.; Project administration: A.M.-A.; Funding acquisition: A.M.-A.

