## Supplementary Material for "Sfp1 integrates TORC1 and PKA activity towards yeast ribosome biogenesis"

#### Supplementary Tables

##### Supplementary Table 1.

| Plasmid | Insert | Modifications | Source |
|---|---|---|---|
| pLV57 | HO5'-SFP1p-**Sfp1**-Linker[1]-pHtdGFP-CYC1t-KanMx-HO3' | - | This study |
| pLV58 | HO5'-SFP1p-***sfp1*$^{\mathbf{TOR7A}}$**-Linker-pHtdGFP-CYC1t-KanMx-HO3' | S39A, S170A, S181A, S183A, T227A, S228A, T446A | This study |
| pLV59 | HO5'-SFP1p-***sfp1*$^{\mathbf{S105A}}$**-Linker-pHtdGFP-CYC1t-KanMx-HO3' | S105A | This study |
| pLV60 | HO5'-SFP1p-***sfp1*$^{\mathbf{S136A}}$**-Linker-pHtdGFP-CYC1t-KanMx-HO3' | S136A | This study |
| pLV3 | HO5'-SFP1p-***sfp1*$^{\mathbf{PKA2A}}$**-Linker-pHtdGFP-CYC1t-KanMx-HO3' | S105A, S136A | This study |
| pLV6 | HO5'-SFP1p-***sfp1*$^{\mathbf{PKA2D}}$**-Linker-pHtdGFP-CYC1t-KanMx-HO3' | S105D, S136D | This study |
| pLV4 | HO5'-SFP1p-***sfp1*$^{\mathbf{TOR7A-PKA2D}}$**-Linker-pHtdGFP-CYC1t-KanMx-HO3' | S39A, S105D, S136D, S170A, S181A, S183A, T227A, S228A, T446A | This study |
| pLV8 | HO5'-SFP1p-***sfp1*$^{\mathbf{TOR7A-PKA2A}}$**-Linker-pHtdGFP-CYC1t-KanMx-HO3' | S39A, S105A, S136A, S170A, S181A, S183A, T227A, S228A, T446A | This study |
| pLV9 | HO5'-SFP1p-***sfp1*$^{\mathbf{TOR7D-PKA2D}}$**-Linker-pHtdGFP-CYC1t-KanMx-HO3' | S39D, S105D, S136D, S170D, S181D, S183D, T227D, S228D, T446D | This study |

| | | | |
|---|---|---|---|
| pLV30 | HO5'-SFP1p-***sfp1***$^{\text{Zn1}}$-Linker-pHtdGFP-CYC1t-KanMx-HO3' | C605A, H618A, H623A | This study |
| pLV31 | HO5'-SFP1p-***sfp1***$^{\text{Zn2}}$-Linker-pHtdGFP-CYC1t-KanMx-HO3' | C661A, H664A, H677A, H680A | This study |
| pLV26 | HO5'-SFP1p-***sfp1***$^{\text{Zn3}}$-Linker-pHtdGFP-CYC1t-KanMx-HO3' | C605A, H618A, H623A, C661A, H664A, H677A, H680A | This study |
| pLV10 | HO5'-SFP1p-***sfp1***$^{\text{TOR7D-PKA2D\_Zn1}}$-Linker-pHtdGFP-CYC1t-KanMx-HO3' | S39D, S105D, S136D, S170D, S181D, S183D, T227D, S228D, T446D, C605A, H618A, H623A | This study |
| pLV11 | HO5'-SFP1p-***sfp1***$^{\text{TOR7D-PKA2D\_Zn2}}$-Linker-pHtdGFP-CYC1t-KanMx-HO3' | S39D, S105D, S136D, S170D, S181D, S183D, T227D, S228D, T446D, C661A, H664A, H677A, H680A | This study |
| pLV32 | HO5'-SFP1p-***sfp1***$^{\text{TOR7D-PKA2D\_Zn3}}$-Linker-pHtdGFP-CYC1t-KanMx-HO3' | S39D, S105D, S136D, S170D, S181D, S183D, T227D, S228D, T446D, C605A, H618A, H623A, C661A, H664A, H677A, H680A | This study |
| pLV33 | HO5'-SFP1p-***sfp1***$^{\Delta\text{Cter}}$-Linker-pHtdGFP-CYC1t-KanMx-HO3' | Δ490-683 | This study |
| pLV34 | HO5'-SFP1p-***sfp1***$^{\Delta\text{Nter}}$-Linker-pHtdGFP-CYC1t-KanMx-HO3' | Δ1-202 | This study |
| pLV61 | HO5'-SFP1p-***sfp1***$^{\Delta\text{NES}}$-Linker-pHtdGFP-CYC1t-KanMx-HO3' | Δ98-106 | This study |
| pLV62 | HO5'-SFP1p-***sfp1***$^{\text{StrongNES}}$-Linker-pHtdGFP-CYC1t-KanMx-HO3' | A99L | This study |
| pLV49 | HO5'-SFP1p-***sfp1***$^{\text{13D}}$-Linker-pHtdGFP-CYC1t-KanMx-HO3' | S39D, S105D, S136D, S170D, S172D, S181D, S183D, S209D, T227D, S228D, S401D, T446D, S646D | This study |
| pLV54 | HO5'-SFP1p-***sfp1***$^{\text{13D\_Zn3}}$-Linker-pHtdGFP-CYC1t-KanMx-HO3' | S39D, S105D, S136D, S170D, S172D, S181D, S183D, S209D, T227D, S228D, S401D, T446D, C605A, H618A, H623A, S646D, C661A, H664A, H677A, H680A | This study |
| pLV50 | HO5'-SFP1p-***sfp1***$^{\text{13A}}$-Linker-pHtdGFP-CYC1t-KanMx-HO3' | S39A, S105A, S136A, S170A, S172A, S181A, S183A, S209A, T227A, S228A, S401A, T446A, S646A | This study |

[1] The linker sequence can be found in the materials and methods section.

Supplementary Table 2.

| Strain | Genotype | Source |
|---|---|---|
| YSBN6 | MATa FY3 HO::HphMX4 | Canelas, 2010 |
| YSBN6 Hta2-mRFP | YSBN6, Hta2::Hta2-mRFP-Ble | This study |
| YSBN6 Sfp1 Hta2-mRFP | YSBN6 Hta2-mRFP, SFP1::SFP1p-**Sfp1**-Linker-pHtdGFP-CYC1t-NatMX4 | Guerra, 2022 |
| YSBN6 TPK1-3AS Hta2-mRFP | YSBN6 Hta2-mRFP, Tpk1-M164G, Tpk2-M147G, Tpk3-M165G | Guerra, 2022 |
| HL215 | W303-1A, sfp1-1::URA3::HIS3 | Lempiäinen, 2009 |
| HL350 | W303-1A, sfp1-Zn2-3A::URA3::HIS3 | Lempiäinen, 2009 |
| HL351 | W303-1A, sfp1-Zn3-4A::URA3::HIS3 | Lempiäinen, 2009 |
| YLV86 | YSBN6 Hta2-mRFP, HO::SFP1p-**Sfp1**-Linker-pHtdGFP-CYC1t-KanMX4 | This study |
| YLV76 | YSBN6 Hta2-mRFP, HO::SFP1p-***sfp1***$^{\mathbf{TOR7A}}$-Linker-pHtdGFP-CYC1t-KanMX4 | This study |
| YLV77 | YSBN6 Hta2-mRFP, HO::SFP1p-***sfp1***$^{\mathbf{S105A}}$-Linker-pHtdGFP-CYC1t-KanMX4 | This study |
| YLV78 | YSBN6 Hta2-mRFP, HO::SFP1p-***sfp1***$^{\mathbf{S136A}}$-Linker-pHtdGFP-CYC1t-KanMX4 | This study |
| YLV10 | YSBN6 Hta2-mRFP, HO::SFP1p-***sfp1***$^{\mathbf{PKA2A}}$-Linker-pHtdGFP-CYC1t-KanMX4 | This study |
| YLV12 | YSBN6 Hta2-mRFP, HO::SFP1p-***sfp1***$^{\mathbf{PKA2D}}$-Linker-pHtdGFP-CYC1t-KanMX4 | This study |
| YLV11 | YSBN6 Hta2-mRFP, HO::SFP1p-***sfp1***$^{\mathbf{TOR7A\text{-}PKA2D}}$-Linker-pHtdGFP-CYC1t-KanMX4 | This study |
| YLV14 | YSBN6 Hta2-mRFP, HO::SFP1p-***sfp1***$^{\mathbf{TOR7A\text{-}PKA2A}}$-Linker-pHtdGFP-CYC1t-KanMX4 | This study |
| YLV16 | YSBN6 Hta2-mRFP, HO::SFP1p-***sfp1***$^{\mathbf{TOR7D\text{-}PKA2D}}$-Linker-pHtdGFP-CYC1t-KanMX4 | This study |
| YLV79 | YSBN6 Hta2-mRFP, HO::SFP1p-***sfp1***$^{\mathbf{Zn1}}$-Linker-pHtdGFP-CYC1t-KanMX4 | This study |
| YLV80 | YSBN6 Hta2-mRFP, HO::SFP1p-***sfp1***$^{\mathbf{Zn2}}$-Linker-pHtdGFP-CYC1t-KanMX4 | This study |

| | | |
|---|---|---|
| YLV38 | YSBN6 Hta2-mRFP, HO::SFP1p-***sfp1***$^{Zn3}$-Linker-pHtdGFP-CYC1t-KanMX4 | This study |
| YLV27 | YSBN6 Hta2-mRFP, HO::SFP1p-***sfp1***$^{TOR7D-PKA2D-Zn1}$-Linker-pHtdGFP-CYC1t-KanMX4 | This study |
| YLV28 | YSBN6 Hta2-mRFP, HO::SFP1p-***sfp1***$^{TOR7D-PKA2D-Zn2}$-Linker-pHtdGFP-CYC1t-KanMX4 | This study |
| YLV37 | YSBN6 Hta2-mRFP, HO::SFP1p-***sfp1***$^{TOR7D-PKA2D-Zn3}$-Linker-pHtdGFP-CYC1t-KanMX4 | This study |
| YLV81 | YSBN6 Hta2-mRFP, HO::SFP1p-***sfp1***$^{\Delta Cter}$-Linker-pHtdGFP-CYC1t-KanMX4 | This study |
| YLV82 | YSBN6 Hta2-mRFP, HO::SFP1p-***sfp1***$^{\Delta Nter}$-Linker-pHtdGFP-CYC1t-KanMX4 | This study |
| YLV83 | YSBN6 Hta2-mRFP, HO::SFP1p-***sfp1***$^{\Delta NES}$-Linker-pHtdGFP-CYC1t-KanMX4 | This study |
| YLV84 | YSBN6 Hta2-mRFP, HO::SFP1p-***sfp1***$^{StrongNES}$-Linker-pHtdGFP-CYC1t-KanMX4 | This study |
| YLV64 | YSBN6 Hta2-mRFP, HO::SFP1p-***sfp1***$^{13D}$-Linker-pHtdGFP-CYC1t-KanMX4 | This study |
| YLV72 | YSBN6 Hta2-mRFP, HO::SFP1p-***sfp1***$^{13D-Zn3}$-Linker-pHtdGFP-CYC1t-KanMX4 | This study |
| YLV65 | YSBN6 Hta2-mRFP, HO::SFP1p-***sfp1***$^{13A}$-Linker-pHtdGFP-CYC1t-KanMX4 | This study |
| YLV87 | YSBN6 TPK1-3AS Hta2-mRFP, HO::SFP1p-***sfp1***-Linker-pHtdGFP-CYC1t-KanMX4 | This study |
| YLV23 | YSBN6 TPK1-3AS Hta2-mRFP, HO::SFP1p-***sfp1***$^{TOR7A}$-Linker-pHtdGFP-CYC1t-KanMX4 | This study |
| YLV19 | YSBN6 TPK1-3AS Hta2-mRFP, HO::SFP1p-***sfp1***$^{PKA2A}$-Linker-pHtdGFP-CYC1t-KanMX4 | This study |
| YLV20 | YSBN6 TPK1-3AS Hta2-mRFP, HO::SFP1p-***sfp1***$^{PKA2D}$-Linker-pHtdGFP-CYC1t-KanMX4 | This study |
| YLV24 | YSBN6 TPK1-3AS Hta2-mRFP, HO::SFP1p-***sfp1***$^{TOR7D-PKA2D}$-Linker-pHtdGFP-CYC1t-KanMX4 | This study |
| YLV22 | YSBN6 TPK1-3AS Hta2-mRFP, HO::SFP1p-***sfp1***$^{TOR7A-PKA2D}$-Linker-pHtdGFP-CYC1t-KanMX4 | This study |
| YLV69 | YSBN6 TPK1-3AS Hta2-mRFP, HO::SFP1p-***sfp1***$^{13D}$-Linker-pHtdGFP-CYC1t-KanMX4 | This study |
| YLV71 | YSBN6 TPK1-3AS Hta2-mRFP, HO::SFP1p-***sfp1***$^{13D-Zn3}$-Linker-pHtdGFP-CYC1t-KanMX4 | This study |

| | | |
|---|---|---|
| YLV70 | YSBN6 TPK1-3AS Hta2-mRFP, HO::SFP1p-***sfp1***$^{\mathbf{13A}}$-Linker-pHtdGFP-CYC1t-KanMX4 | This study |
| YLV41 | YSBN6 TPK1-3AS Hta2-mRFP, HO::SFP1p-***sfp1***$^{\mathbf{Zn3}}$-Linker-pHtdGFP-CYC1t-KanMX4 | This study |

### Supplementary Figures

#### Supplementary Figure 1.

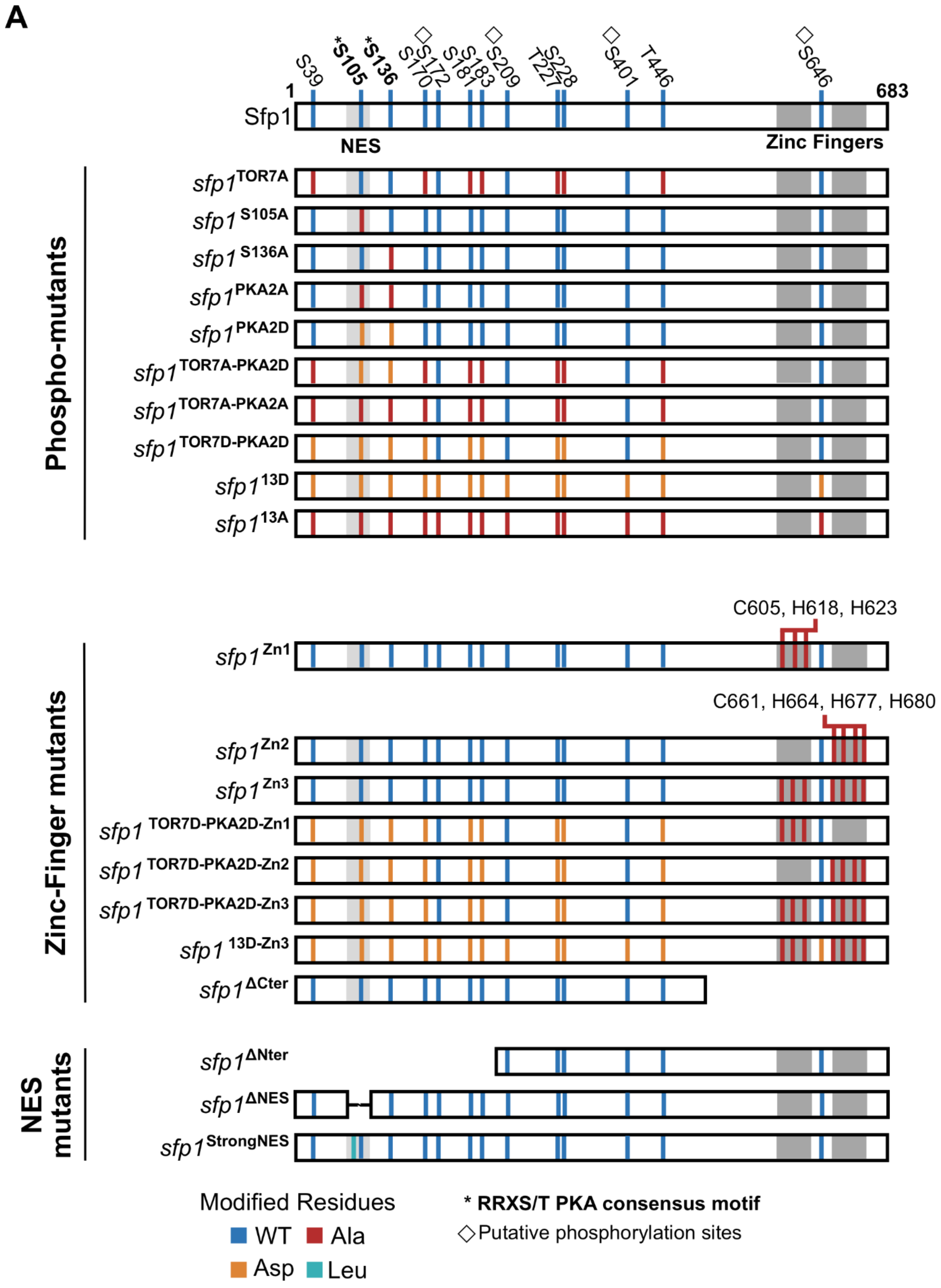

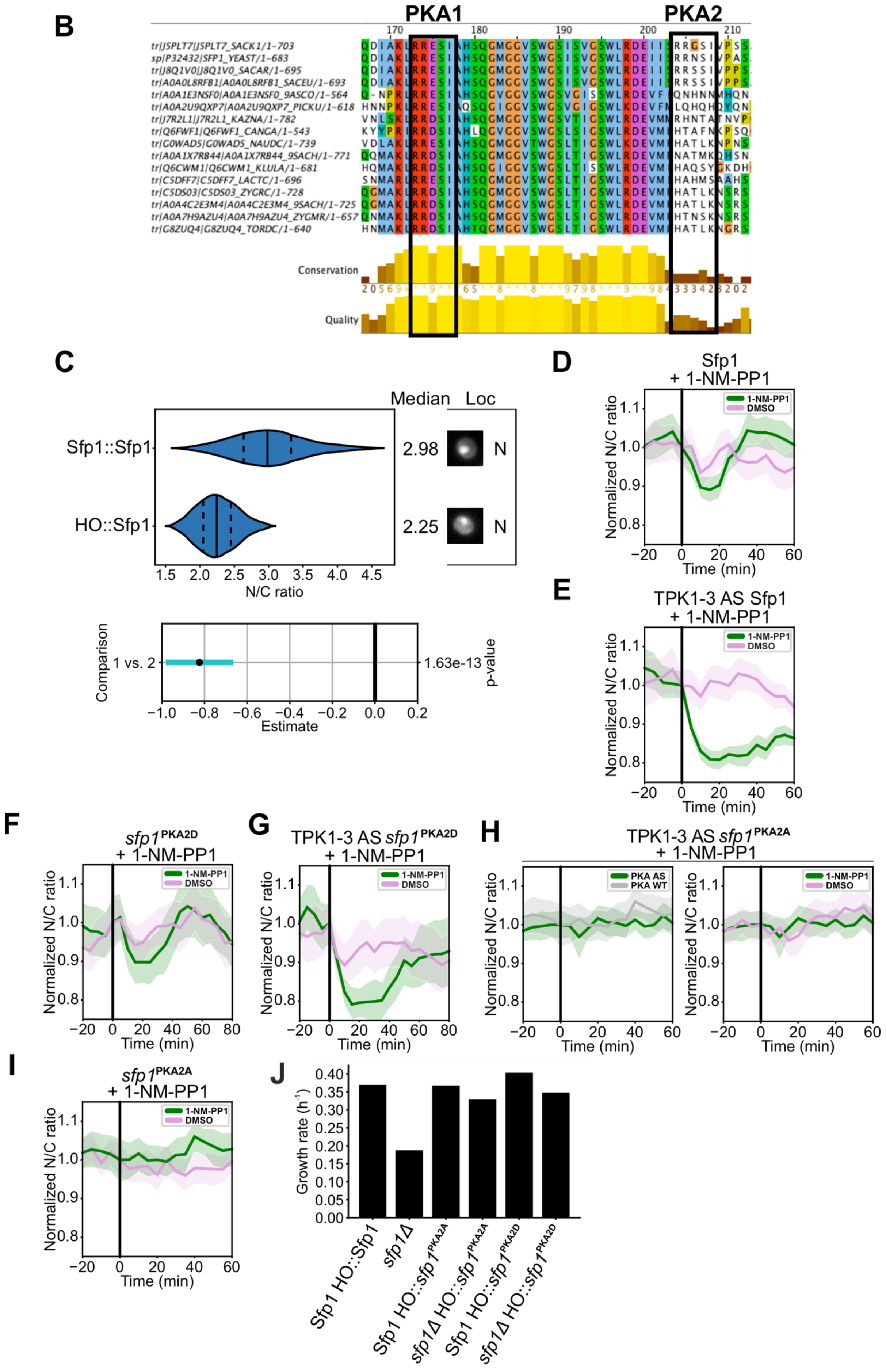
B
PKA1
PKA2
170
180
190
200
210
tr|J5PLT7|J5PLT7_SACK1/1-703
sp|P32432|SFP1_YEAST/1-683
tr|J8Q1V0|J8Q1V0_SACAR/1-695
tr|A0A0L8RFB1|A0A0L8RFB1_SACEU/1-693
tr|A0A1E3NSF0|A0A1E3NSF0_9ASCO/1-564
tr|A0A2U9QXP7|A0A2U9QXP7_PICKU/1-618
tr|J7R2L1|J7R2L1_KAZNA/1-782
tr|Q6FWF1|Q6FWF1_CANGA/1-543
tr|G0WAD5|G0WAD5_NAUDC/1-739
tr|A0A1X7RB44|A0A1X7RB44_9SACH/1-771
tr|Q6CWM1|Q6CWM1_KLULA/1-681
tr|C5DFF7|C5DFF7_LACTC/1-696
tr|C5DS03|C5DS03_ZYGRC/1-728
tr|A0A4C2E3M4|A0A4C2E3M4_9SACH/1-725
tr|A0A7H9AZU4|A0A7H9AZU4_ZYGMR/1-657
tr|G8ZUQ4|G8ZUQ4_TORDC/1-640
Conservation
Quality
C
Median
Loc
Sfp1::Sfp1
HO::Sfp1
2.98
2.25
N
N
1.5 2.0 2.5 3.0 3.5 4.0 4.5
N/C ratio
Comparison
1 vs. 2
1.63e-13
p-value
−1.0 −0.8 −0.6 −0.4 −0.2 0.0 0.2
Estimate
D
Sfp1
+ 1-NM-PP1
1-NM-PP1
DMSO
Normalized N/C ratio
Time (min)
E
TPK1-3 AS Sfp1
+ 1-NM-PP1
F
sfp1PKA2D
+ 1-NM-PP1
G
TPK1-3 AS sfp1PKA2D
+ 1-NM-PP1
H
TPK1-3 AS sfp1PKA2A
+ 1-NM-PP1
PKA AS
PKA WT
I
sfp1PKA2A
+ 1-NM-PP1
J
Growth rate (h-1)
0.00 0.05 0.10 0.15 0.20 0.25 0.30 0.35 0.40
Sfp1 HO::Sfp1
sfp1Δ
Sfp1 HO::sfp1PKA2A
sfp1Δ HO::sfp1PKA2A
Sfp1 HO::sfp1PKA2D
sfp1Δ HO::sfp1PKA2D

(**A**) Schematic representation of the collection of all yeast strains expressing wild-type and mutant Sfp1-pHtdGFP fusions used in this study. The TORC1 phosphorylation sites (blue marks, normal font), C-terminal Zn finger domains (dark grey segments), putative leucine-rich type NES region (light grey segment), the region containing the putative PKA phosphorylation sites (star mark, bold) and putative phosphorylation sites not linked to a kinase (blue marks, diamond shape) are indicated. (**B**) Jalview visualization of a Multiple sequence alignment (MUSCLE algorithm, cluster method: neighbour joining) of 21 Sfp1 and Sfp1-like proteins across several yeast genera. PKA1 and PKA2 are denoted (black rectangles). Protein accession numbers, sequence conservation and quality of the alignment are displayed. (**C**) Sfp1 nuclear to cytosolic (N/C) ratio distributions in cells expressing Sfp1 in the endogenous Sfp1 locus (Sfp1::Sfp1; n = 39) and cells expressing Sfp1 in the HO locus as a second copy (HO::Sfp1; n = 39). Median (solid line) and 25th and 75th percentiles (dashed lines) are displayed. Representative cells images are shown on the right of the violin plots. Statistical comparison of means was carried out with the Games-Howell test. The p-value of the comparison is indicated alongside the corresponding 99% confidence interval of the mean difference. The significantly different distribution is indicated with a green confidence interval. Legend: 1 = Sfp1::Sfp1, 2 = HO::Sfp1.(**D**) Sfp1 N/C ratio dynamics in response to 1-NM-PP1 addition in wild-type cells treated with 1-NM-PP1 (n = 37) compared to cells treated with the drug vehicle DMSO (n = 39). Off-target 1-NM-PP1 effects induce a transient drop in wild-type Sfp1 nuclear localization while DMSO causes a minor drop. (**E**) Sfp1 N/C ratio dynamics in response to 1-NM-PP1 addition in wild-type cells carrying the analog-sensitive PKA mutations (PKAas) treated with 1-NM-PP1 (n = 50) compared to cells treated with the drug vehicle DMSO (n = 43). Inhibition of PKAas leads to a sustained drop in Sfp1 nuclear localization while DMSO causes a minor drop. (**F**) *sfp1*$^{PKA2D}$ N/C ratio dynamics in response to 1-NM-PP1 addition in wild-type cells treated with 1-NM-PP1 (n = 30) compared to cells treated with the drug vehicle DMSO (n = 37). Off-target 1-NM-PP1 effects induce a transient drop in *sfp1*$^{PKA2D}$ nuclear localization while DMSO causes a minor transient drop. (**G**) *sfp1*$^{PKA2D}$ N/C ratio dynamics in response to 1-NM-PP1 addition in cells carrying the analog-sensitive PKA mutations treated with 1-NM-PP1 (n = 27) compared to cells treated with the drug vehicle DMSO (n = 26). Inhibition of PKAas leads to a transient drop comparable to the 1-NM-PP1 off-target effect observed in wild-type *sfp1*$^{PKA2D}$ while DMSO induces a minor transient drop. The plot extends to 80 min post-treatment to demonstrate the convergence of the control and treated cells. Single-cell measurements of cells carrying the analog-sensitive PKA mutations at 65 min post-perturbation were discarded due to a transient shift in focus. Measurements at 60- and 70-min post-perturbation were linearly interpolated to generate the data point at 65 min. (**H**) *sfp1*$^{PKA2A}$ N/C ratio dynamics in response to 1-NM-PP1 addition in cells carrying the analog-sensitive PKA mutations (n = 42) compared to wild-type treated cells (n = 31) and PKA analog-sensitive cells treated with the drug vehicle DMSO (n = 38). Inhibition of PKAas does not lead to a drop in the N/C ratio of PKAas and wild-type *sfp1*$^{PKA2A}$ cells, and DMSO treatment has no effect either. (**I**) *sfp1*$^{PKA2A}$ N/C ratio dynamics in response to 1-NM-PP1 addition in wild-type cells treated with 1-NM-PP1 (n = 31) compared to cells treated with the drug vehicle DMSO (n = 42). Inhibition of PKAas or treatment with DMSO do not induce a drop in the *sfp1*$^{PKA2A}$ N/C ratio. (**J**) Growth rate of Sfp1 wild type (HO::Sfp1 extra copy), *sfp1Δ*, Sfp1 HO::*sfp1*$^{PKA2A}$, *sfp1Δ sfp1*$^{PKA2A}$, Sfp1 HO::*sfp1*$^{PKA2D}$ and *sfp1Δ sfp1*$^{PKA2D}$ strains.

### Supplementary Figure 2.

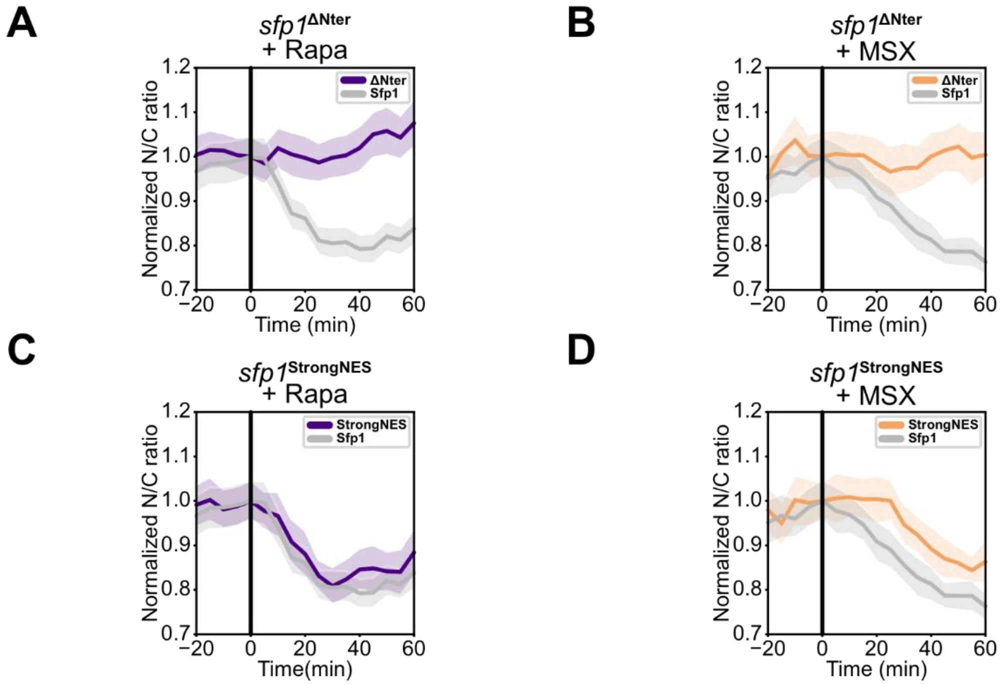


(**A**) Sfp1 N/C ratio dynamics in response to rapamycin in cells carrying wild-type Sfp1 and the *sfp1*$^{\Delta\text{Nter}}$ mutant. Rapamycin leads to a sustained drop in wild-type Sfp1 nuclear localization (n = 47), whereas the *sfp1*$^{\Delta\text{Nter}}$ mutant (n = 75) does not show a drop in *sfp1*$^{\Delta\text{Nter}}$ nuclear localization. (**B**) Sfp1 N/C ratio dynamics in response to methionine sulfoximine in cells carrying wild-type Sfp1 or the *sfp1*$^{\Delta\text{Nter}}$ mutant. Methionine sulfoximine leads to a sustained drop in wild-type Sfp1 nuclear localization (n = 42), whereas the *sfp1*$^{\Delta\text{Nter}}$ mutant (n = 32) does not show a drop in *sfp1*$^{\Delta\text{Nter}}$ nuclear localization. (**C**) Sfp1 N/C ratio dynamics in response to rapamycin in cells carrying wild-type Sfp1 and the *sfp1*$^{\text{StrongNES}}$ mutant. Rapamycin leads to a comparable sustained drop in wild-type Sfp1 nuclear localization (n = 47) and in *sfp1*$^{\text{StrongNES}}$ nuclear localization (n = 23). (**D**) Sfp1 N/C ratio dynamics in response to methionine sulfoximine in cells carrying wild-type Sfp1 or the *sfp1*$^{\Delta\text{StrongNES}}$ mutant. Methionine sulfoximine leads to a comparable sustained drop in wild-type Sfp1 nuclear localization (n = 42) and in *sfp1*$^{\text{StrongNES}}$ nuclear localization (n = 29).

Supplementary Figure 3.

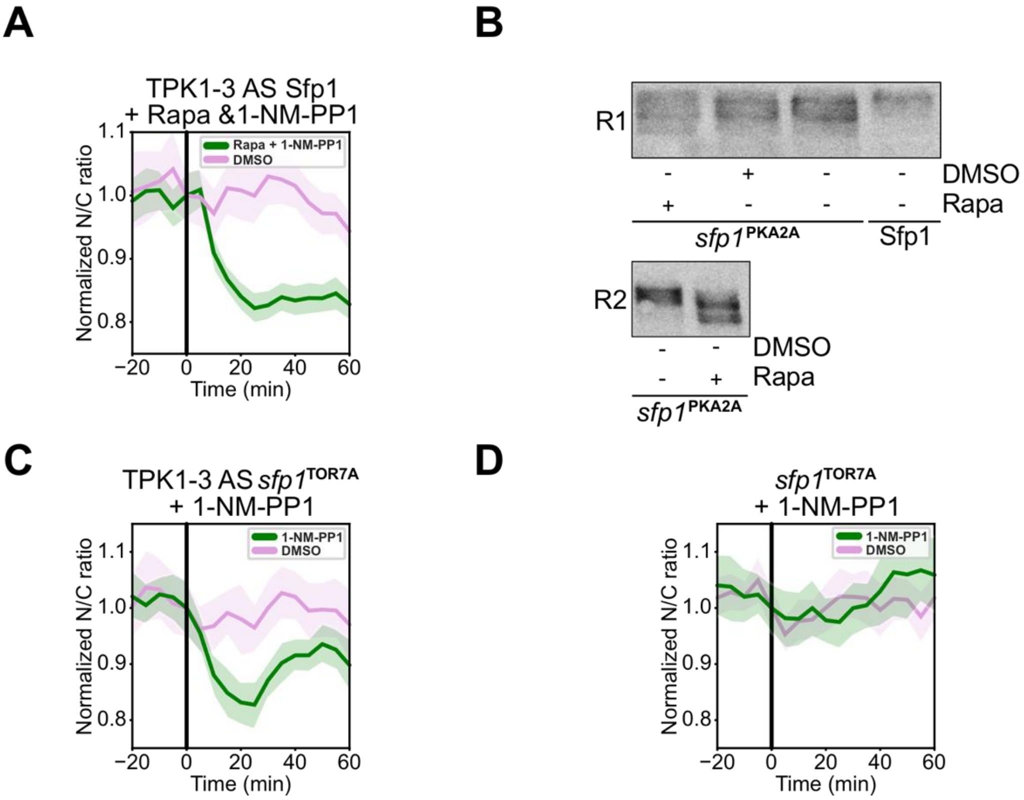


(**A**) Response of Sfp1 to TORC1 and PKA inhibition by a joint treatment with rapamycin and 1-NM-PP1 (n = 52) and the drug vehicle DMSO (n = 43) in the PKAas background. While the joint treatment leads to a sustained drop in Sfp1 nuclear localization of PKA analog-sensitive cells, DMSO only generates a minor drop. (**B**) Phos-tag analyses of the GFP-tagged *sfp1*$^{PKA2A}$ mutant. Rapamycin leads to further decrease in phosphorylation of *sfp1*$^{PKA2A}$ mutant whereas DMSO (drug vehicle) does not cause a decrease. R1 and R2 denote biological replicates. (**C**) *sfp1*$^{TOR7A}$ N/C ratio dynamics in response to 1-NM-PP1 addition in cells carrying the analog-sensitive PKA mutations (n = 24) compared to treatment with the drug vehicle DMSO (n = 23). Inhibition of PKAas leads to a sustained drop in analog-sensitive PKA *sfp1*$^{TOR7A}$ nuclear localization while DMSO causes a minor drop. (**D**) *sfp1*$^{TOR7A}$ N/C ratio dynamics in response to 1-NM-PP1 addition in wild-type cells treated with 1-NM-PP1 (n = 22) compared to a treatment with the drug vehicle DMSO (n = 37). Inhibition of PKAas or treatment with DMSO lead to a minor transient drop in *sfp1*$^{TOR7A}$ nuclear localization.

Supplementary Figure 4.

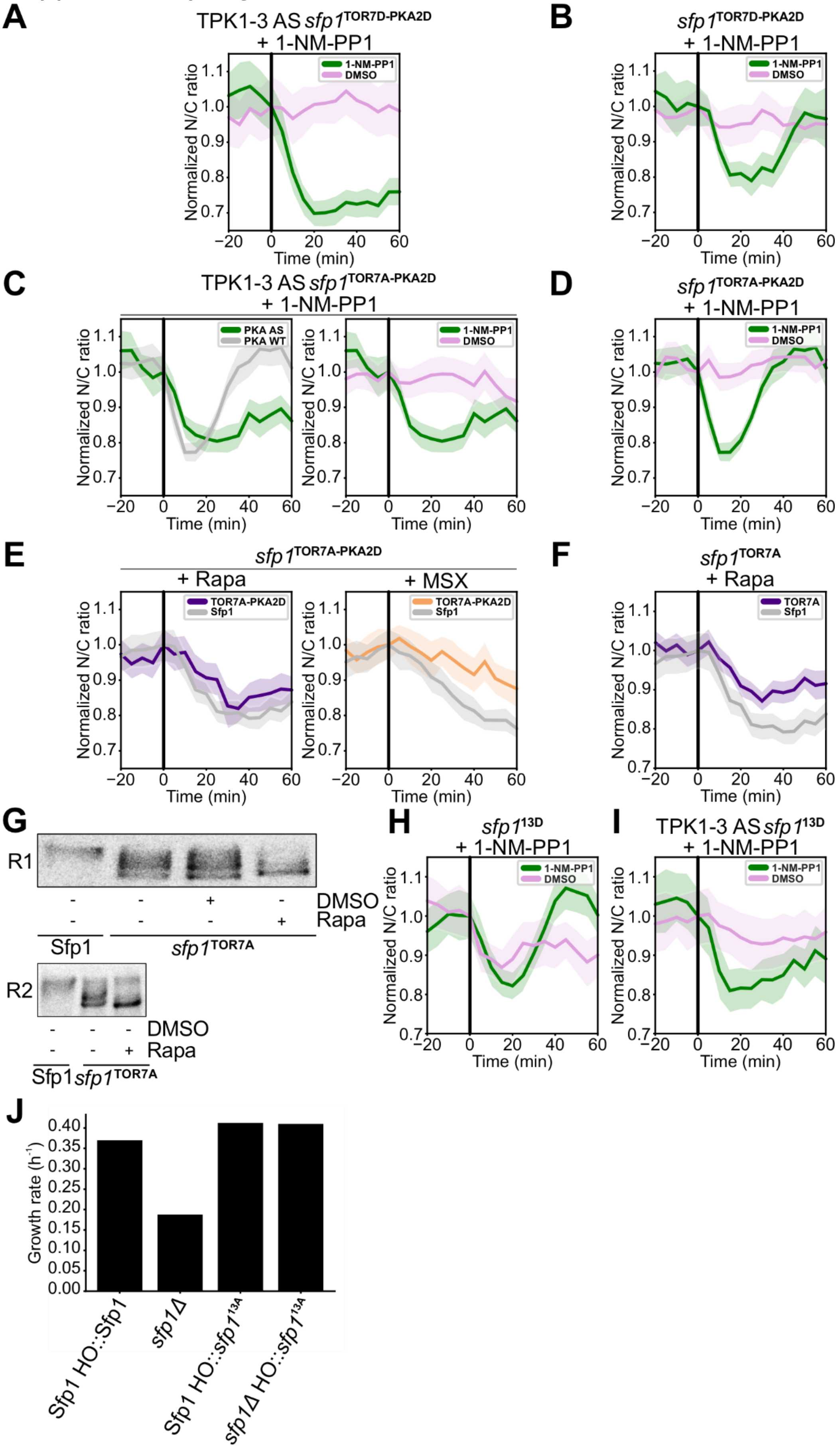

(**A**) *sfp1*$^{\text{TOR7D-PKA2D}}$ N/C ratio dynamics in response to 1-NM-PP1 addition in cells carrying the analog-sensitive PKA mutations (n = 36) compared to a treatment with the drug vehicle DMSO (n = 36). Inhibition of PKAas leads to a sustained drop in analog-sensitive PKA *sfp1*$^{\text{TOR7D-PKA2D}}$ nuclear localization while DMSO does not cause a drop. (**B**) *sfp1*$^{\text{TOR7D-PKA2D}}$ N/C ratio dynamics in response to 1-NM-PP1 addition in wild-type cells (n = 30) compared to treatment with the drug vehicle DMSO (n = 36). Off-target 1-NM-PP1 effects induce a transient drop in *sfp1*$^{\text{TOR7D-PKA2D}}$ nuclear localization while DMSO causes a minor drop. (**C**) *sfp1*$^{\text{TOR7A-PKA2D}}$ N/C ratio dynamics in response to 1-NM-PP1 addition in cells carrying the analog-sensitive PKA mutations (n = 30) compared to wild-type treated cells (n = 40) and PKA analog-sensitive cells treated with the drug vehicle DMSO (n = 41). Inhibition of PKAas leads to a sustained drop in analog-sensitive PKA *sfp1*$^{\text{TOR7A-PKA2D}}$ nuclear localization while off-target 1-NM-PP1 effects induce a transient drop in *sfp1*$^{\text{TOR7A-PKA2D}}$ and DMSO causes a minor a drop. (**D**) *sfp1*$^{\text{TOR7A-PKA2D}}$ N/C ratio dynamics in response to 1-NM-PP1 addition in wild-type cells (n = 40) compared to a treatment with the drug vehicle DMSO (n = 41). Off-target 1-NM-PP1 effects induce a transient drop in *sfp1*$^{\text{TOR7A-PKA2D}}$ nuclear localization while DMSO causes a minor transient drop. (**E**) *sfp1*$^{\text{TOR7A-PKA2D}}$ N/C ratio dynamics in response to rapamycin or methionine sulfoximine in cells carrying wild-type Sfp1 or the *sfp1*$^{\text{TOR7A-PKA2D}}$ mutant. Rapamycin and methionine sulfoximine lead to a sustained drop in *sfp1*$^{\text{TOR7A-PKA2D}}$ nuclear localization (Rapa n = 40; MSX n = 38) comparable to the drop observed in Sfp1 wild-type cells (Rapa n = 47; MSX n = 42). (**F**) *sfp1*$^{\text{TOR7A}}$ N/C ratio dynamics in response to rapamycin in cells carrying wild-type Sfp1 and the *sfp1*$^{\text{TOR7A}}$ mutant. Rapamycin leads to a slightly reduced sustained drop in *sfp1* $^{\text{TOR7A}}$ nuclear localization (n = 71) compared to the drop observed in Sfp1 wild-type cells (n = 47). (**G**) Phos-tag analyses of the GFP-tagged *sfp1*$^{\text{TOR7A}}$ mutant. Rapamycin leads to further decrease in phosphorylation of *sfp1*$^{\text{TOR7A}}$ mutant whereas DMSO (Drug vehicle) does not cause a decrease. R1 and R2 denotes biological replicates. (**H**) *sfp1*$^{\text{13D}}$ N/C ratio dynamics in response to 1-NM-PP1 addition in wild-type cells (n = 24) compared to a treatment with the drug vehicle DMSO (n = 38). Off-target 1-NM-PP1 effects induce a transient drop in *sfp1*$^{\text{13D}}$ nuclear localization while DMSO causes a minor transient drop. (**I**) *sfp1*$^{\text{13D}}$ N/C ratio dynamics in response to 1-NM-PP1 addition in cells carrying the analog-sensitive PKA mutations (n = 27) compared to a treatment with the drug vehicle DMSO (n = 23). Inhibition of PKAas leads to a sustained drop in analog-sensitive PKA *sfp1*$^{\text{13D}}$ nuclear localization while DMSO causes a minor a drop. (**J**) Growth rate of Sfp1 wild type (HO::Sfp1 extra copy), *sfp1Δ*, Sfp1 HO::*sfp1*$^{\text{13A}}$, *sfp1Δ* HO::*sfp1*$^{\text{13A}}$ strains.

Supplementary Figure 5.

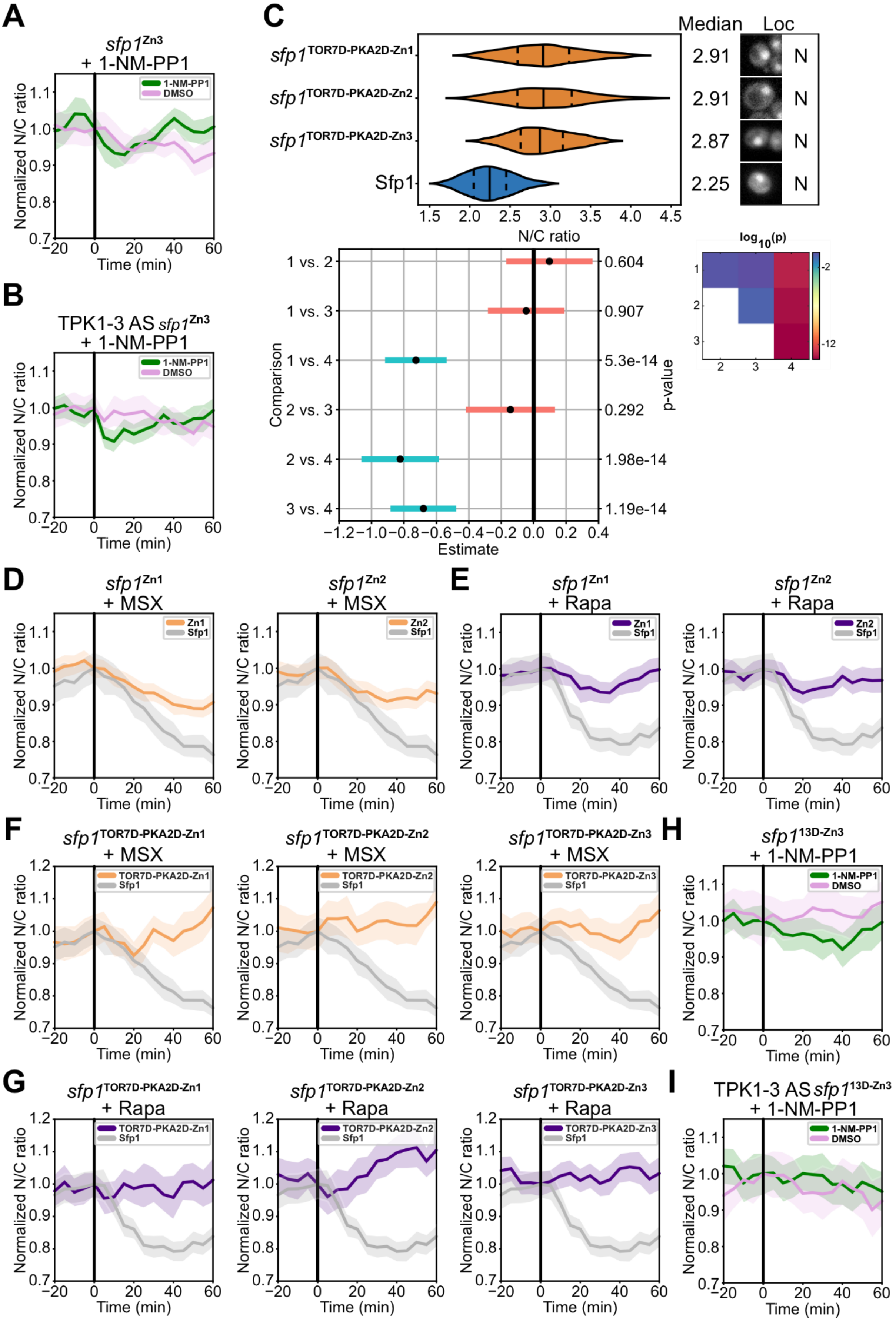

(**A**) *sfp1*$^{\text{Zn3}}$ N/C ratio dynamics in response to 1-NM-PP1 addition in wild-type cells (n = 37) compared to a treatment with the drug vehicle DMSO (n = 43). Off-target 1-NM-PP1 effects induce a transient drop in *sfp1*$^{\text{Zn3}}$ nuclear localization while DMSO causes a minor drop. (**B**) *sfp1*$^{\text{Zn3}}$ N/C ratio dynamics in response to 1-NM-PP1 addition in cells carrying the analog-sensitive PKA mutations (n = 46) compared to a treatment with the drug vehicle DMSO (n = 36). Inhibition of PKAas leads to a transient drop comparable to the 1-NM-PP1 off-target effect observed in wild-type *sfp1*$^{\text{Zn3}}$ while DMSO induces a minor drop. (**C**) Sfp1 nuclear to cytosolic (N/C) ratio distributions in cells carrying the *sfp1*$^{\text{TOR7D-PKA2D-Zn1}}$ (n = 58), *sfp1*$^{\text{TOR7D-PKA2D-Zn2}}$ (n = 32), *sfp1*$^{\text{TOR7D-PKA2D-Zn3}}$ (n = 66) mutants and Sfp1 wild-type (n = 39). Median (solid line) and 25th and 75th percentiles (dashed lines) are displayed. Representative cells images are shown on the right of the violin plots. Statistical comparison of means was carried out with the Games-Howell test. P-values of all pairwise comparisons are graphically summarized in a matrix and indicated alongside the corresponding 99% confidence intervals of all mean differences. Not significantly different distributions are indicated with red confidence intervals while significantly different distributions are indicated with green confidence intervals. Legend: 1 = *sfp1*$^{\text{TOR7D-PKA2D-Zn1}}$, 2 = *sfp1*$^{\text{TOR7D-PKA2D-Zn2}}$, 3 = *sfp1*$^{\text{TOR7D-PKA2D-Zn3}}$, 4 = Sfp1. (**D**) *sfp1*$^{\text{Zn1}}$ & *sfp1*$^{\text{Zn2}}$ N/C ratio dynamics in response to methionine sulfoximine in cells carrying wild-type Sfp1 or the *sfp1*$^{\text{Zn1}}$ & *sfp1*$^{\text{Zn2}}$ mutants. Methionine sulfoximine leads to a reduced drop in *sfp1*$^{\text{Zn1}}$ & *sfp1*$^{\text{Zn2}}$ nuclear localization (*sfp1*$^{\text{Zn1}}$, n = 65; *sfp1*$^{\text{Zn2}}$, n = 63) whereas a larger drop of Sfp1 nuclear localization is observed in Sfp1 wild-type cells (n = 42). (**E**) *sfp1*$^{\text{Zn1}}$ & *sfp1*$^{\text{Zn2}}$ N/C ratio dynamics in response to rapamycin in cells carrying wild-type Sfp1 or the *sfp1*$^{\text{Zn1}}$ & *sfp1*$^{\text{Zn2}}$ mutants. Rapamycin leads to a reduced drop in *sfp1*$^{\text{Zn1}}$ & *sfp1*$^{\text{Zn2}}$ nuclear localization (*sfp1*$^{\text{Zn1}}$, n = 68; *sfp1*$^{\text{Zn2}}$, n = 61) whereas a larger drop of Sfp1 nuclear localization is observed in Sfp1 wild-type cells (n = 47). (**F**) *sfp1*$^{\text{TOR7A-PKA2D-Zn1}}$, *sfp1*$^{\text{TOR7A-PKA2D-Zn2}}$ & *sfp1*$^{\text{TOR7A-PKA2D-Zn3}}$ N/C ratio dynamics in response to methionine sulfoximine in cells carrying wild-type Sfp1 or the *sfp1*$^{\text{TOR7A-PKA2D-Zn1}}$, *sfp1*$^{\text{TOR7A-PKA2D-Zn2}}$ & *sfp1*$^{\text{TOR7A-PKA2D-Zn3}}$ mutants. Methionine sulfoximine does not lead to a drop in *sfp1*$^{\text{TOR7A-PKA2D-Zn1}}$, *sfp1*$^{\text{TOR7A-PKA2D-Zn2}}$ & *sfp1*$^{\text{TOR7A-PKA2D-Zn3}}$ nuclear localization (*sfp1*$^{\text{TOR7A-PKA2D-Zn1}}$, n= 38; *sfp1*$^{\text{TOR7A-PKA2D-Zn2}}$, n = 25; *sfp1*$^{\text{TOR7A-PKA2D-Zn3}}$, n = 41) whereas a drop of Sfp1 nuclear localization is observed in Sfp1 wild-type cells (n = 42). (**G**) *sfp1*$^{\text{TOR7A-PKA2D-Zn1}}$, *sfp1*$^{\text{TOR7A-PKA2D-Zn2}}$ & *sfp1*$^{\text{TOR7A-PKA2D-Zn3}}$ N/C ratio dynamics in response to rapamycin in cells carrying wild-type Sfp1 or the *sfp1*$^{\text{TOR7A-PKA2D-Zn1}}$, *sfp1*$^{\text{TOR7A-PKA2D-Zn2}}$ & *sfp1*$^{\text{TOR7A-PKA2D-Zn3}}$ mutants. Rapamycin does not lead to a drop in *sfp1*$^{\text{TOR7A-PKA2D-Zn1}}$, *sfp1*$^{\text{TOR7A-PKA2D-Zn2}}$ & *sfp1*$^{\text{TOR7A-PKA2D-Zn3}}$ nuclear localization (*sfp1*$^{\text{TOR7A-PKA2D-Zn1}}$, n= 30; *sfp1*$^{\text{TOR7A-PKA2D-Zn2}}$, n = 34; *sfp1*$^{\text{TOR7A-PKA2D-Zn3}}$, n = 48) whereas a drop of Sfp1 nuclear localization is observed in Sfp1 wild-type cells (n = 47). (**H**) *sfp1*$^{\text{13D-Zn3}}$ N/C ratio dynamics in response to 1-NM-PP1 addition in wild-type cells (n = 35) compared to treatment with the drug vehicle DMSO (n = 29). Off-target 1-NM-PP1 effects induce a minor transient drop in *sfp1*$^{\text{13D-Zn3}}$ nuclear localization while DMSO does not cause a drop. (**I**) *sfp1*$^{\text{13D-Zn3}}$ N/C ratio dynamics in response to 1-NM-PP1 addition in cells carrying the analog-sensitive PKA mutations (n = 37) compared to a treatment with the drug vehicle DMSO (n = 30). Inhibition of PKAas leads to a minor drop comparable to the 1-NM-PP1 off-target effect observed in wild-type *sfp1*$^{\text{13D-Zn3}}$ while DMSO induces a minor drop.

### Supplementary Figure 6.

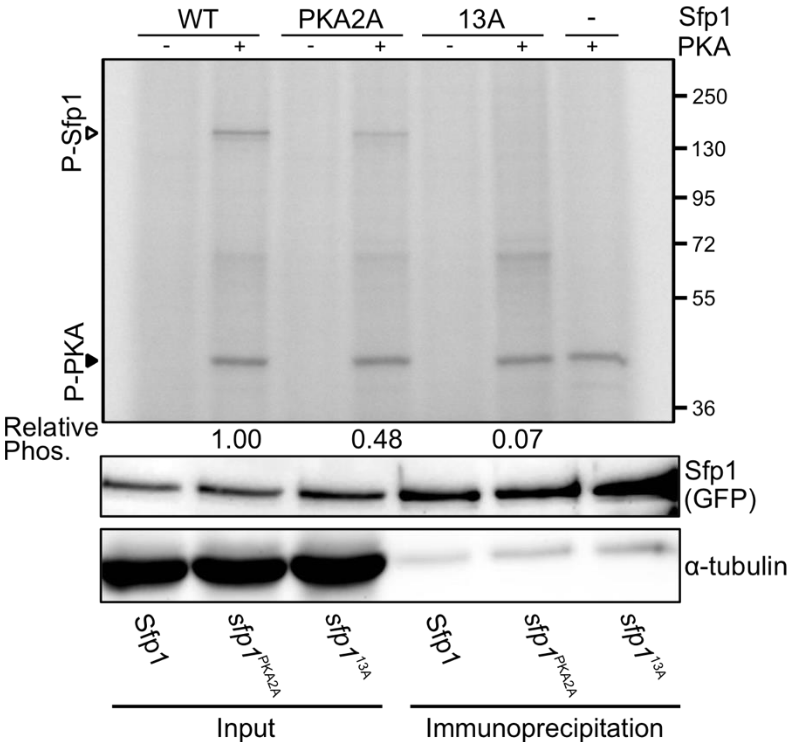


Phosphorylation of Sfp1 by protein kinase A: following the treatment of PKAas cells with rapamycin and 1-NM-PP1 (to remove endogenous Sfp1 phosphorylation), Sfp1-pHtdGFP, *sfp1*$^{PKA2A}$-pHtdGFP and *sfp1*$^{13A}$-pHtdGFP were immunoprecipitated by GFP-Trap (lower panel), and were incubated with γ-$^{32}$P-ATP, with or without bovine PKA for 30 min at 30°C (Upper panel). The reaction mixtures were separated by SDS-PAGE, the gel was exposed to storage phosphor screen, and autoradiography was performed by Cyclone plus (Perkin Elmer). The open triangle indicates the phosphorylated Sfp1 and the closed triangle indicates the (auto)phosphorylated PKA. Relative phosphorylation of Sfp1 in the three strains (Materials and Methods) is indicated below the autoradiograph.

Supplementary Figure 7.

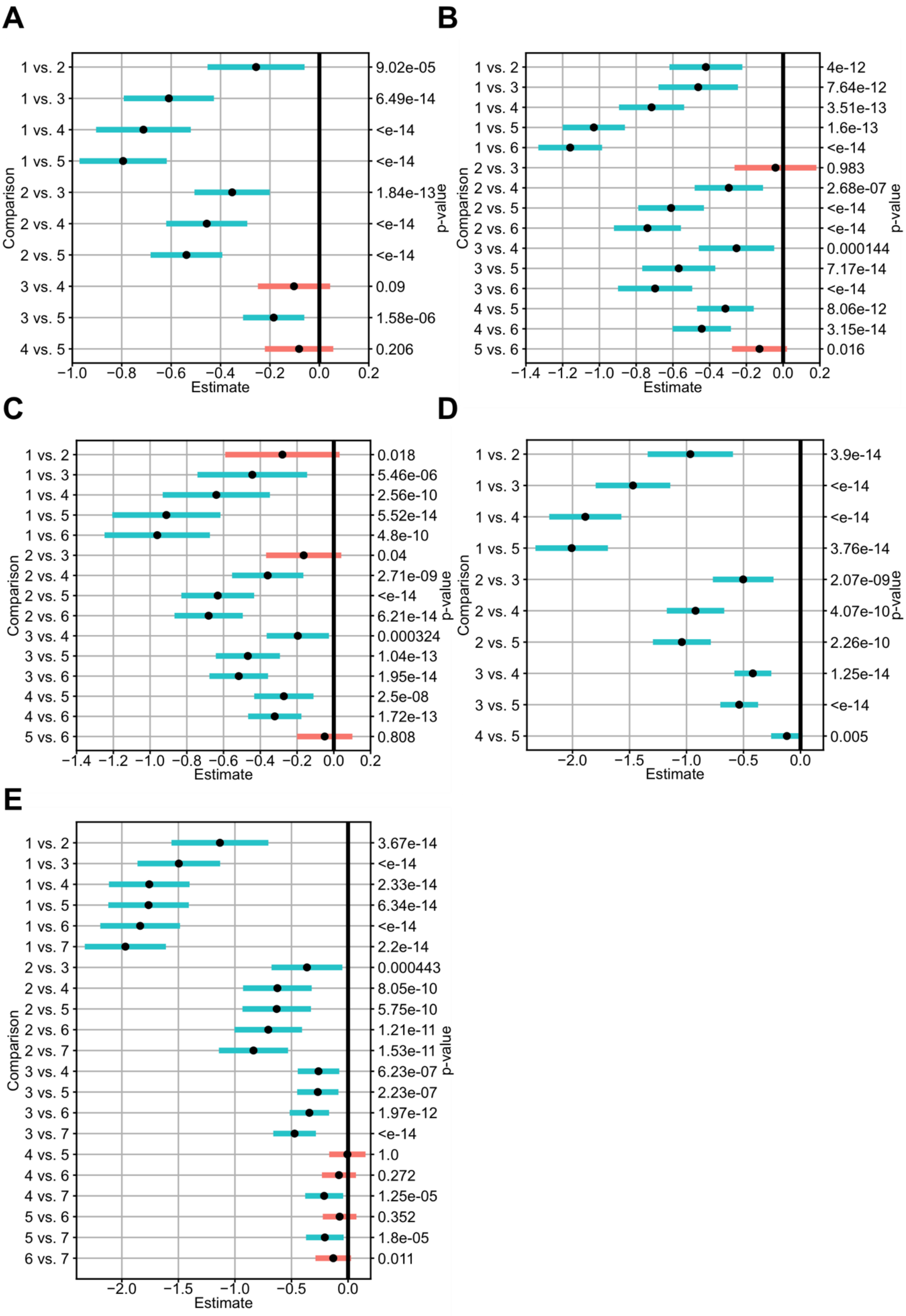

Statistical comparisons of the Sfp1 (and mutants) Nuclear to Cytosolic (N/C) ratio distributions carried out with the Games-Howell test. P-values associated with each pairwise comparison is indicated alongside the corresponding 99% confidence interval of the mean difference. Not significantly different distributions are indicated with red confidence intervals while significantly different distributions are indicated with green confidence intervals. (**A**) Refers to Fig.1C. Legend: 1 = $sfp1^{PKA2D}$, 2 = Sfp1, 3 = $sfp1^{S136A}$, 4 = $sfp1^{PKA2A}$, 5 = $sfp1^{S105A}$. (**B**) Refers to Fig.2C. Legend: 1 = $sfp1^{\Delta Nter}$, 2 = $sfp1^{\Delta NES}$, 3 = $sfp1^{PKA2D}$, 4 = Sfp1, 5 = $sfp1^{StrongNES}$, 6 = $sfp1^{PKA2A}$. (**C**) Refers to Fig.3D. Legend: 1 = $sfp1^{TOR7D\text{-}PKA2D}$, 2 = $sfp1^{TOR7A\text{-}PKA2D}$, 3 = Sfp1, 4 = $sfp1^{TOR7A}$, 5 = $sfp1^{PKA2A}$, 6 = $sfp1^{TOR7A\text{-}PKA2A}$. (**D**) Refers to Fig.4C. Legend: 1 = $sfp1^{13D}$, 2 = $sfp1^{TOR7D\text{-}PKA2D}$, 3 = Sfp1, 4 = $sfp1^{13A}$, 5 = $sfp1^{TOR7A\text{-}PKA2A}$. (**E**) Refers to Fig.4C. Legend: 1 = $sfp1^{13D}$, 2 = $sfp1^{13D\text{-}Zn3}$, 3 = Sfp1, 4 = $sfp1^{Zn3}$, 5 = $sfp1^{Zn2}$, 6 = $sfp1^{Zn1}$, 7 = $sfp1^{\Delta Cter}$.
